# MULTIPLE, REDUNDANT CARBOXYLIC ACID TRANSPORTERS SUPPORT MITOCHONDRIAL METABOLISM IN *PLASMODIUM FALCIPARUM*

**DOI:** 10.1101/2024.11.26.624872

**Authors:** Krithika Rajaram, Gabriel W. Rangel, Justin T. Munro, Sethu C. Nair, Manuel Llinás, Sean T. Prigge

## Abstract

The mitochondrion of the deadliest human malaria parasite, *Plasmodium falciparum,* is an essential source of cellular acetyl-CoA during the asexual blood-stage of the parasite life cycle. Blocking mitochondrial acetyl-CoA synthesis leads to a hypoacetylated proteome and parasite death. We previously determined that mitochondrial acetyl-CoA is primarily synthesized from glucose-derived pyruvate by α-ketoacid dehydrogenases. Here, we asked if inhibiting the import of glycolytic pyruvate across the mitochondrial inner membrane would affect acetyl-CoA production and, thus, could be a potential target for antimalarial drug development. We selected the two predicted mitochondrial pyruvate carrier proteins (*Pf*MPC1 and *Pf*MPC2) for genetic knockout and isotopic metabolite tracing via HPLC-MS metabolomic analysis. Surprisingly, we observed that asexual blood-stage parasites could survive the loss of either or both *Pf*MPCs with only minor growth defects, despite a substantial reduction in the amount of glucose-derived isotopic labelling into acetyl-CoA. Furthermore, genetic deletion of two additional mitochondrial carboxylic acid transporters – DTC (di/tricarboxylic acid carrier) and YHM2 (a putative citrate/α-ketoglutarate carrier protein) – only mildly affected asexual blood-stage replication, even in the context of *Pf*MPC deficiency. Although we observed no added impact on the incorporation of glucose carbon into acetyl-CoA in these quadruple knockout mutants, we noted a large decrease in glutamine-derived label in tricarboxylic acid cycle metabolites, suggesting that DTC and YHM2 both import glutamine derivatives into the mitochondrion. Altogether, our results expose redundant routes used to fuel the blood-stage malaria parasite mitochondrion with imported carbon from two major sources – glucose and glutamine.

**SIGNIFICANCE:** The mitochondrion of malaria parasites generates key molecules, such as acetyl-CoA, that are required for numerous cellular processes. To support mitochondrial biosynthetic pathways, the parasites must transport carbon sources into this organelle. By studying how the mitochondrion obtains pyruvate, a molecule derived from glucose, we have uncovered redundant carbon transport systems that ensure parasite survival in red blood cells. This metabolic redundancy poses a challenge for drug development, as it enables the parasite to adapt and survive by relying on alternative pathways when one is disrupted.

## INTRODUCTION

*Plasmodium falciparum* is a protozoan parasite that causes the deadliest form of malaria in humans, killing over 600,000 people annually (World Health Organization, 2023). Each malaria parasite contains a single mitochondrion that is required throughout its life cycle. Despite its essentiality, only a few critical functions have been attributed to the mitochondrion during the symptomatic stage of disease when parasites largely replicate asexually within human red blood cells. The tricarboxylic acid (TCA) cycle is dispensable in the asexual blood stage (ABS) forms (Ke et al., 2015; Rajaram et al., 2022), and the primary purpose of the mitochondrial electron transport chain (mETC) is to furnish ubiquinone for dihydroorotate dehydrogenase (DHODH), a mitochondrial enzyme that participates in *de novo* pyrimidine synthesis (Painter et al., 2007). As a result, nearly all mitochondrial-specific antimalarials have been developed to target either the mETC or DHODH. However, in sexual blood-stage parasites, other mitochondrial enzymes are critical to development including aconitase (Ke et al., 2015) and isocitrate dehydrogenase (Yang et al., 2022). To expand the druggable mitochondrial proteome, the organelle has been the focus of intense research to further our understanding of its biology and metabolic output.

In line with these efforts, we recently established that the *P. falciparum* mitochondrion is responsible for generating cellular acetyl-CoA in ABS parasites (Nair et al., 2023). This production is largely driven by mitochondrial pyruvate dehydrogenase (mPDH), also known as the branched chain alpha-keto acid dehydrogenase. The multisubunit mPDH complex uses glucose-derived carbon, likely in the form of pyruvate, a three-carbon carboxylic acid product of glycolysis, as its substrate. Although a second PDH complex exists in the apicoplast organelle in *Plasmodium*, it is dispensable in ABS parasites (Cobbold et al., 2013; Pei et al., 2010; Swift et al., 2020a). When mPDH is disrupted, parasites become reliant on the TCA cycle enzyme α-ketoglutarate dehydrogenase (KDH) for survival. KDH is an α-ketoacid dehydrogenase that is closely related to mPDH and that primarily converts α-ketoglutarate into succinyl-CoA in parasites; however, *in vitro*, it has also been demonstrated to catalyze the conversion of pyruvate to acetyl-CoA (Chan et al., 2013). The two enzymes constitute a synthetic lethal pair, and their inactivation results in the hypoacetylation of histones and other proteins outside the mitochondrion (Nair et al., 2023).

A recent study has shown that 5-15% of the cellular acetate pool in mammalian cells is derived from ROS-mediated pyruvate decarboxylation (Liu et al., 2018). While non-enzymatic conversion of pyruvate to acetate may also occur in *Plasmodium* parasites, it is insufficient to sustain parasites lacking functional mPDH and KDH, which instead require supplementation with millimolar quantities of acetate for survival (Nair et al., 2023). A nuclear acetyl-CoA synthetase (ACS) is responsible for scavenging and ligating free acetate to CoA (Nair et al., 2023; Prata et al., 2021; Summers et al., 2021). However, given the high concentration of acetate required for this bypass to be effective, ACS is unlikely to be a significant generator of acetyl-CoA under physiological conditions (Nair et al., 2023). Combined with recent observations that pyruvate and mitochondrial energy metabolism are linked to artemisinin resistance, the acetyl-CoA biosynthetic pathway has emerged as a promising target for drug discovery (Chen et al., 2014; de Vries et al., 2022; Mok et al., 2021; Schalkwijk et al., 2019; Zhang et al., 2022).

Pyruvate is generated from the breakdown of glucose during glycolysis and must be imported from the cytoplasm into the mitochondrial matrix for use by mPDH/KDH (Nair et al., 2023). In many eukaryotes, the transport of ions, metabolites and proteins across the outer mitochondrial membrane is mediated by a highly abundant porin called the voltage-dependent anion channel (VAC) (Hodge and Colombini, 1997; Homblé et al., 2012; Mallo et al., 2021; Wideman et al., 2013). *P. falciparum* has a putative VAC that was recently demonstrated to be essential for parasite survival (Anaguano et al., 2023). The protein localizes primarily to a membrane at the parasite-host interface but also appears to overlap partially with the mitochondrion. The inner mitochondrial membrane is far more selective, and entry is typically mediated by specific solute transporters (Cunningham and Rutter, 2020). *P. falciparum* encodes two proteins that are homologous to the mitochondrial pyruvate carrier proteins MPC1 and MPC2 found in yeast, mammals, and the related apicomplexan parasite *Toxoplasma gondii* (Bricker et al., 2012; Herzig et al., 2012; Lyu et al., 2023; Weiner and Kooij, 2016; Wunderlich, 2022). These are small proteins (12 kDa and 15 kDa, respectively) that form part of a heteromeric complex of about 150 kDa in other organisms (Bricker et al., 2012; Herzig et al., 2012).

In this study, we probed whether the putative MPC proteins in *P. falciparum* (hereafter referred to as *Pf*MPC) play a role in acetyl-CoA synthesis by transporting pyruvate into the mitochondrion. We employed CRISPR-based gene deletion methods to discover that both proteins are in fact dispensable in ABS parasites. However, there is a marked decrease in the incorporation of ^13^C glucose into acetyl-CoA and its derivatives in *Pf*MPC-deficient parasites, confirming the role of the *Pf*MPCs in pyruvate import. Deletion of two additional carboxylic acid transporters, a mitochondrial di/tricarboxylate carrier (DTC) and a putative citrate/α-ketoglutarate transporter (YHM2), in the parasite line lacking the two *Pf*MPCs did not further impact survival or ^13^C glucose incorporation into known mitochondrial metabolites but did impact ^13^C glutamine incorporation into TCA cycle metabolites. Overall, our data indicate that malaria parasites employ redundant mitochondrial carboxylate transporters to fuel the TCA cycle with carbon derived from glucose and glutamine.

## RESULTS

### Mitochondrial carrier proteins *Pf*MPC1 and *Pf*MPC2 are dispensable for blood-stage growth of *P. falciparum*

Pyruvate is conventionally imported into mitochondria for conversion to acetyl-CoA. This process is mediated by members of the mitochondrial pyruvate carrier (MPC) family. While three paralogs of this family have been found in yeast, human and *Plasmodium* genomes only encode two MPC proteins (Bricker et al., 2012; Herzig et al., 2012; Wunderlich, 2022). *Pf*MPC1 and *Pf*MPC2 have been assigned strong localization scores by two different computational methods (MitoProtII and PlasmoMitoCarta) (Claros and Vincens, 1996; van Esveld et al., 2021). To confirm these predictions, we examined the cellular location of the putative *Pf*MPC1 (PF3D7_1340800) and *Pf*MPC2 (PF3D7_1470400) proteins in *P. falciparum* by generating parasite lines with second copies of *Pf*MPC1 or *Pf*MPC2 fused to a C-terminal SFG (superfolder GFP) tag (Pédelacq et al., 2006; Roberts et al., 2019) (**Fig. S1A-D**). Unfortunately, the *Pf*MPC1-SFG signal was too faint to be measured by live-cell imaging and was undetectable by immunofluorescence microscopy. However, an MPC1 homolog in the closely related parasite *T. gondii* has been experimentally established as a mitochondrial protein (Lyu et al., 2023). Live microscopy analysis of the transgenic parasites showed that *Pf*MPC2-SFG colocalized with the MitoTracker probe, confirming that *Pf*MPC2 is a mitochondrial protein (**Fig. 1A**).

**Fig. 1.**
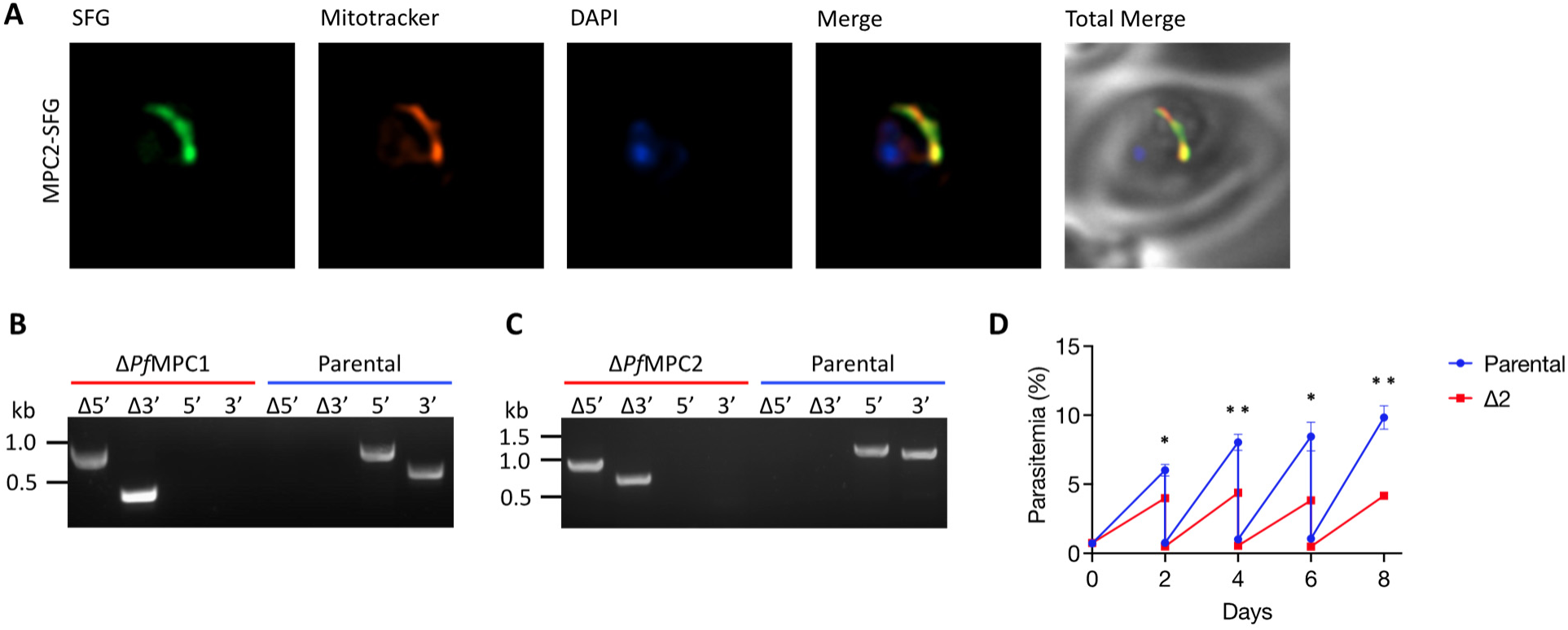
*Pf*MPC proteins are dispensable in ABS parasites. (A) Representative immunofluorescence microscopy images of *Pf*MPC2-SFG parasites show that *Pf*MPC2-SFG (green) colocalizes with MitoTracker (red) [Mander’s coefficient M1 (green in red) = 0.97, SD = 0.015, n = 19]. Images represent fields that are 10 μm long by 10 μm wide. Gene deletion of (B) *Pf*MPC1 and (C) *Pf*MPC2 in NF54^attB^ (parental) parasites was confirmed by PCR amplification (Δ5’ and Δ3’ products). Intact gene loci (5’ and 3’ products) were detected only in the parental (blue) and not in the mutant lines (red). (D) Growth of Δ2 and parental parasites was monitored by flow cytometry over an 8-day period. Cultures were seeded at 0.5% parasitemia on day 0 and diluted 1:8 every other day. Two biological experiments were conducted in quadruplicate (two-way ANOVA computed using Prism 10 [GraphPad Software, Inc], with Šídák’s multiple comparison test; *, p ≤ 0.05; **, p ≤ 0.01). Error bars represent the standard deviations from the mean.

To determine if *Pf*MPC1 and *Pf*MPC2 have essential roles in blood-stage parasites, we used CRISPR/Cas9 to knock out the genes individually in an NF54^attB^ parasite line (Adjalley et al., 2010) (**Fig. S2**). Successful gene deletions were confirmed by diagnostic PCR (**Fig. 1B-C**), indicating that both proteins are dispensable in ABS parasites. In yeast, MPC1 forms a heteromeric complex with MPC2 or MPC3, and deletion of either participating subunit renders the transporter non-functional (Bender et al., 2015; Tavoulari et al., 2019). However, it has recently been found that homotypic complexes of human MPC2 are capable of transporting pyruvate in the absence of MPC1 (Nagampalli et al., 2018). To determine if one MPC protein can substitute for the other in *P. falciparum*, we attempted to create a double-knockout line by targeting *Pf*MPC1 for deletion in Δ*Pf*MPC2 parasites. To provide a possible metabolic bypass, we divided the transfected parasite culture into two flasks; one flask contained media that was supplemented with 5 mM acetate in the event that the deletion of both *Pf*MPC proteins blocked all acetyl-CoA production in the mitochondrion.

To our surprise, we obtained parasites lacking *Pf*MPC1 and *Pf*MPC2 (hereafter referred to as Δ2) in media with or without acetate (**Fig. S3A-C**). We compared the growth of Δ2 parasites against the parental line and observed only a moderate ABS fitness defect (**Fig. 1D**). These results suggest that *Pf*MPC1 and *Pf*MPC2 either do not play a major role in supporting acetyl-CoA synthesis in *P. falciparum*, or that a different mechanism is able to partially compensate for their absence.

### *Pf*MPCs are required for optimal incorporation of glucose carbon into acetyl-CoA

Once pyruvate enters the mitochondrion, it is acted upon by α-ketoacid dehydrogenases to yield the two-carbon-donor compound acetyl-CoA. Acetyl-CoA then enters the TCA cycle by combining with the four-carbon molecule oxaloacetate to form six-carbon citrate. Citrate is converted in two steps to α-ketoglutarate, which also serves as an entry point for carbon derived from glutamine into the TCA cycle (**Fig. 2A**). Our previously published results indicate that acetyl-CoA or an unknown derivative is also exported out of the mitochondrion to acetylate proteins and metabolites in the cytosol and nucleus (Nair et al., 2023). To determine if the levels of acetyl-CoA and downstream products are affected in Δ2 compared to parental parasites, we replaced unlabeled glucose or glutamine in the culture media with isotopically labeled ^13^C_6_-glucose or ^13^C_5_-glutamine, respectively, and traced carbon incorporation into various metabolites using HPLC-MS metabolomics (**Fig. 2A**).

**Fig. 2.**
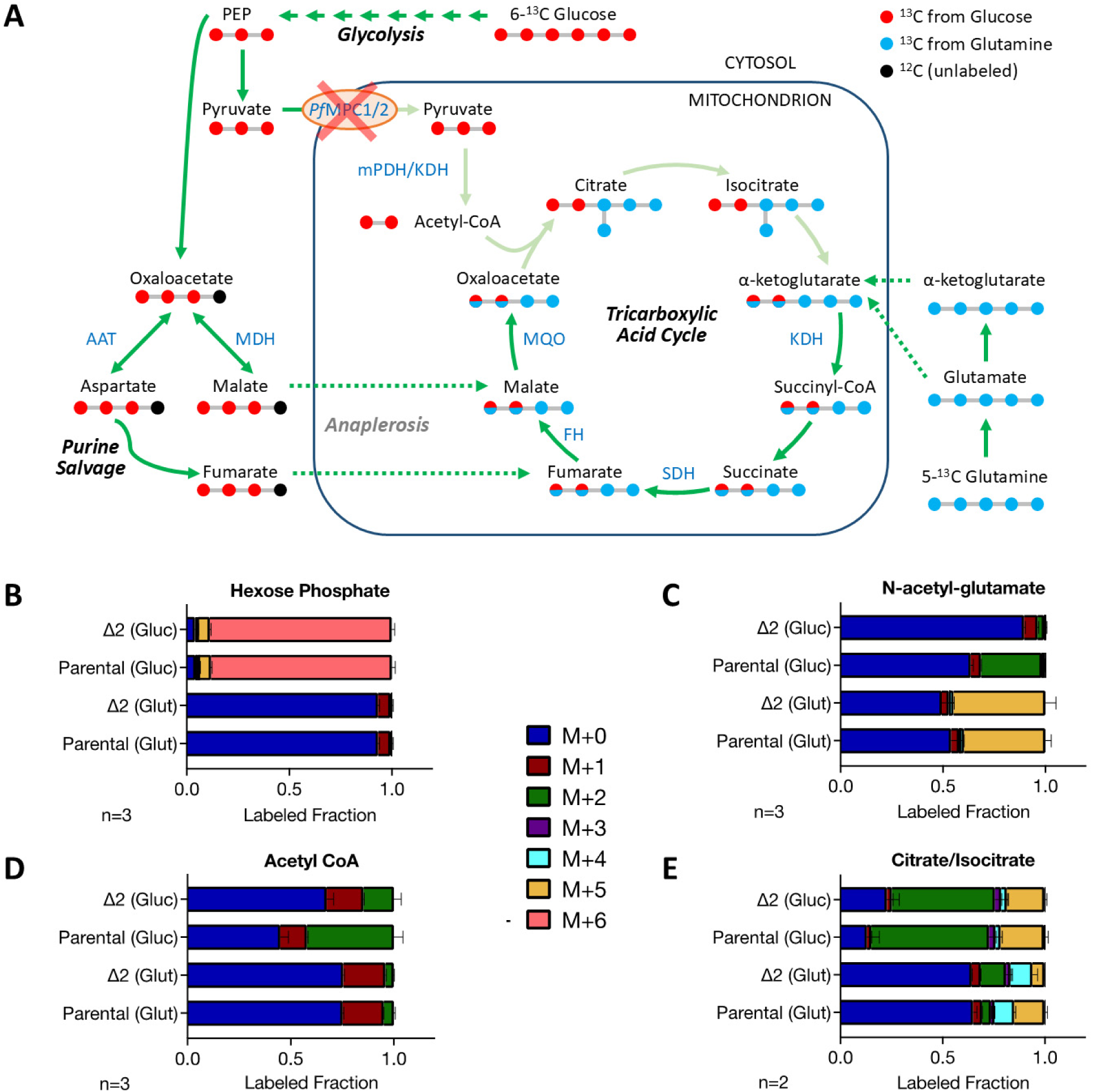
*Pf*MPC deletion reduces the synthesis of acetyl-CoA from glucose. (A) Schematic depicting the labeling pattern of mitochondrial metabolites in Δ2 parasites after short-term incubation with uniformly labeled ^13^C-glucose (solid red circles) or ^13^C-glutamine (solid blue circles). Black circles represent unlabeled atoms. Circles colored with both red and blue indicate carbons that may be derived from either ^13^C-glucose or ^13^C-glutamine. Arrows represent enzymatic reactions and are colored based on the known flow (dark green), reduced flow (light green) or predicted flow (dotted green) of carbon atoms. Abbreviations: PEP – phosphoenolpyruvate; AAT – aspartate aminotransferase; MDH – malate dehydrogenase; mPDH – mitochondrial pyruvate dehydrogenase; KDH – α-ketoglutarate dehydrogenase; SDH – succinate dehydrogenase; FH – fumarate dehydrogenase; MQO – malate-quinone oxidoreductase. (B-E) Fraction of isotopically labeled metabolites when parental or Δ2 parasites are incubated with labeled glucose (Gluc) or glutamine (Glut). Color coding denotes the mass (M) of the parent compound plus the mass shift from the incorporation of heavy carbon isotope (^13^C). Labeling data are expressed as the fraction of the total metabolite pool observed in ‘n’ independent experiments. Error bars represent the standard deviations from the mean.

We found that over 86% of hexose phosphate and over 60% of α-ketoglutarate were derived from ^13^C-glucose and ^13^C-glutamine, respectively, over the course of the 2.5-hour labeling period in both parasite lines (**Fig. 2B**, **Fig. S4A**). As expected, the fraction of labeling from ^13^C-glutamine in TCA cycle intermediates and N-acetyl-glutamate, an acetylated amino acid derived from acetyl-CoA, did not vary significantly between the two lines, ruling out any intrinsic variations in these metabolic pathways between the parental and Δ2 parasites (**Fig. 2C-E**, **Fig. S4**).

We next examined the incorporation of ^13^C label from glucose into acetyl-CoA and N-acetyl-glutamate (**Fig. 2C-D**). In parental parasites, both metabolites were labeled with two carbons from ^13^C_6_-glucose (M+2); by contrast, this labeling was highly reduced, though not absent, in Δ2 parasites. The integrity of this pathway is also evident from the levels of two-carbon and five-carbon (M+5; with three oxaloacetate carbons derived from anaplerotic sources) labeled citrate species, which are only slightly reduced in Δ2 compared to parental parasites (**Fig. 2E**). Similarly, negligible reductions are observed in the M+2 glucose label in downstream TCA cycle metabolites (**Fig. S4**). These results suggest that *Pf*MPC1 and *Pf*MPC2 are the primary mitochondrial pyruvate carriers, and their loss negatively impacts acetyl-CoA synthesis and parasite growth. Despite this, *Pf*MPC deficiency is not lethal likely because some glucose-derived carbon is still available in the mitochondrion to produce sufficient levels of acetyl-CoA.

### DTC and YHM2 are dispensable individually and together with *Pf*MPCs

While *Pf*MPC1 and *Pf*MPC2 appear responsible for the vast majority of pyruvate import into the *P. falciparum* mitochondrion, a limited amount of glucose carbon still enters the organelle in their absence (**Fig. 2**, **Fig. S4**). To determine if other mitochondrial carboxylate transporters can substitute for *Pf*MPCs, we searched the annotated *P. falciparum* genome using resources on PlasmodDB.org for putative mitochondrial carboxylate carriers and found two candidates (Alvarez-Jarreta et al., 2024; Aurrecoechea et al., 2009; Harb et al., 2024). The first, a di/tricarboxylic acid transporter (DTC; PF3D7_0823900), is a known mitochondrial protein that has been shown to exchange several carboxylates in reconstituted liposomes (Nozawa et al., 2011; van Dooren et al., 2006); however, its affinity for pyruvate, a monocarboxylate, has not been evaluated. The second candidate, YHM2 (PF3D7_1223800), a homolog of the yeast citrate/α-ketoglutarate transporter, has not been characterized in *P. falciparum* (Castegna et al., 2010). To determine its subcellular location, we generated transgenic parasites expressing a second copy of SFG-tagged YHM2 (**Fig. S1A-B**, **Fig. S5**) and confirmed the colocalization of YHM2-SFG with MitoTracker by live microscopy (**Fig. 3A**).

**Fig. 3.**
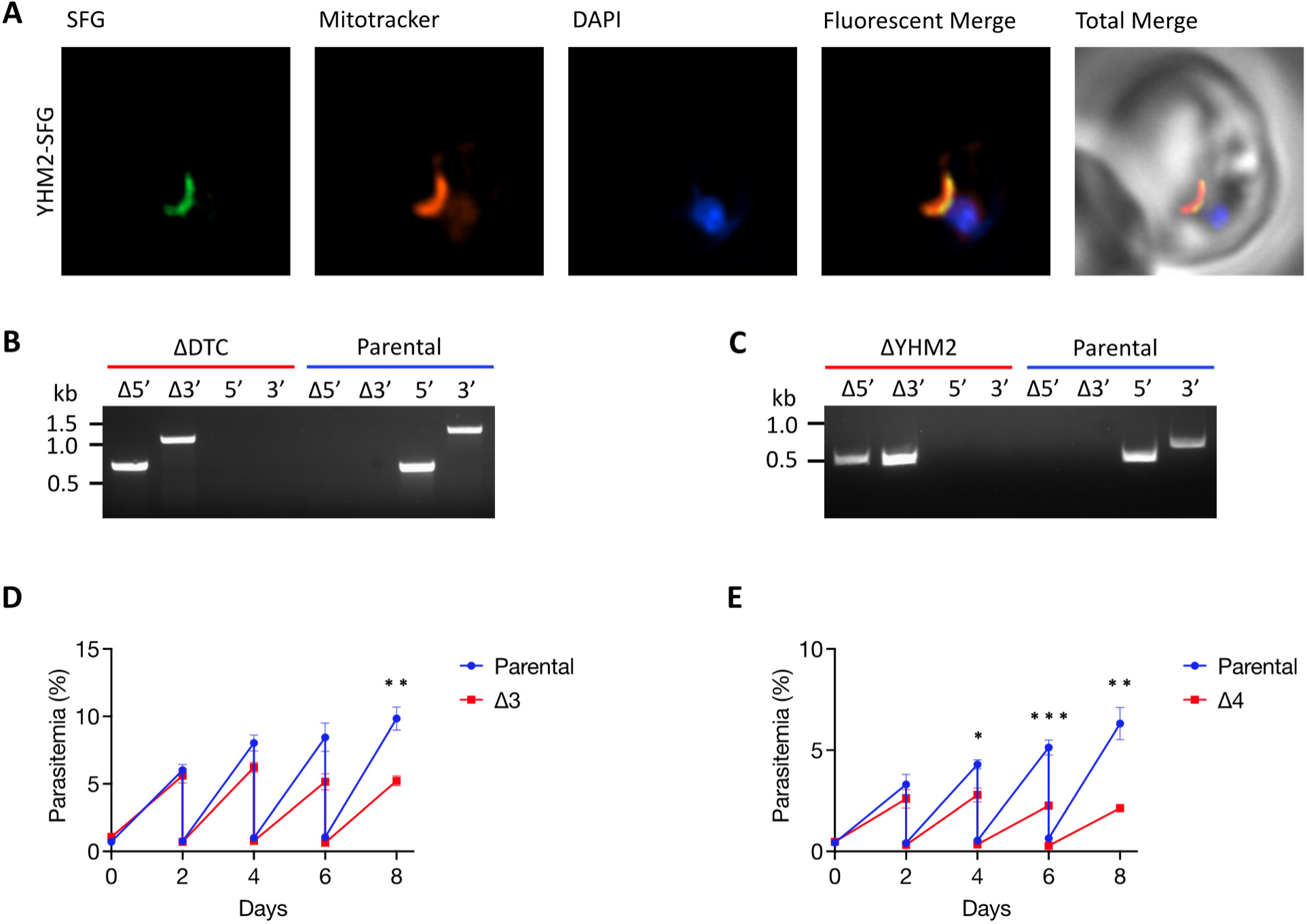
Multiple transporters are dispensable in ABS parasites. (A) Representative immunofluorescence microscopy images of YHM2-SFG parasites show that YHM2-SFG (green) colocalizes with MitoTracker (red) [Mander’s coefficient M1 (green in red) = 0.97, SD = 0.026, n = 16]. Images represent fields that are 10 μm long by 10 μm wide. Gene deletion of (B) DTC and (C) YHM2 in NF54^attB^ (parental) parasites was confirmed by PCR amplification (Δ5’ and Δ3’ products). Intact gene loci (5’ and 3’ products) were detected only in the parental (blue) and not in the mutant lines (red). Growth of parental, (D) Δ3, and (E) Δ4 parasites was compared by measuring their initial and final parasitemia over the 48-h developmental cycle within red blood cells. The average fold change in parasitemia was calculated from the ratio of these values to provide a measure of fitness. The data are from eight biological experiments with quadruplicate samples. Error bars represent standard deviations from the mean. Statistics: Unpaired t-test using Prism 10 [GraphPad Software, Inc]; ns – not significant. Note: The growth data for the Δ3 line was collected alongside Δ2, so the control parental data shown in Fig. 3D is identical to that in Fig. 1D.

To determine the essentiality of DTC and YHM2 in *P. falciparum*, we attempted to delete the genes individually in the parental NF54^attB^ line (**Fig. S2A**). Parasites lacking DTC or YHM2 were confirmed by PCR (**Fig. 3B-C**, **Fig. S6**). Next, we used the same CRISPR plasmids to sequentially disrupt the coding regions of DTC and YHM2 in Δ2 parasites. As before, we split the transfected parasite culture into selective media with and without acetate. Drug-resistant parasites were obtained in both acetate-free and supplemented media. We verified DTC and YHM2 deletions in clonal lines using diagnostic PCR as well as by whole genome sequencing (**Fig. S7**, **Fig. S8**). Growth comparisons of the triple (Δ*Pf*MPC1/*Pf*MPC2/DTC or Δ3) and quadruple (Δ*Pf*MPC1/*Pf*MPC2/DTC/YHM2 or Δ4) mutants with parental parasites revealed only moderate differences in replication rates (**Fig. 3D-E**). These results indicate that DTC and YHM2 are both dispensable in ABS parasites and their absence does not confer large fitness costs to parasites.

### DTC and YHM2 transport α-ketoglutarate

Although DTC and YHM2 are dispensable in ABS parasites, their absence could still affect carbon import into the mitochondrion. To determine if the deletion of these genes in the Δ2 background confers an additive decrease in glucose carbon incorporation into acetyl-CoA, we incubated Δ3, Δ4 and parental parasites in growth media containing ^13^C_6_-glucose. As expected, we observed a significant loss of two-carbon label (M+2) in acetyl-CoA and N-acetyl-glutamate (**Fig. S9B-C**) in Δ3 and Δ4 mutants when compared with parental parasites. However, this reduction was similar to what we observed with the Δ2 parasites (**Fig. 2C-D**). This suggests that DTC and YHM2 do not play a major role in importing glucose carbon into the mitochondrion. As observed with the double mutant, some two- and five-carbon label was still being incorporated into citrate in Δ3 and Δ4 parasites (**Fig. S9D**).

The *P. falciparum* DTC protein has been shown to mediate efficient counter-transport of α-ketoglutarate and other carboxylates *in vitro* (Nozawa et al., 2011). While the substrate specificity of YHM2 has not been determined in malaria parasites, its homolog in yeast is a citrate/α-ketoglutarate antiporter (Castegna et al., 2010). To test if *P. falciparum* DTC and YHM2 import glutamine carbon as α-ketoglutarate to feed the TCA cycle, we supplemented the growth media of Δ3, Δ4, and parental parasites with ^13^C_5_-glutamine. N-acetyl-glutamate and α-ketoglutarate were fully labeled with ^13^C_5_-glutamine carbon to a similar extent in all parasite lines (**Fig. S9C**, **Fig. 4A-B**). On the other hand, we observed a substantial reduction in ^13^C_5_-glutamine-labeled succinate, fumarate, and malate in Δ3 parasites, and this defect was even more pronounced in Δ4 parasites (**Fig. 4C-E**). Taken together, these results suggest that DTC and YHM2 are both responsible for the transport of α-ketoglutarate from the cytosol into the mitochondrion.

**Fig. 4.**
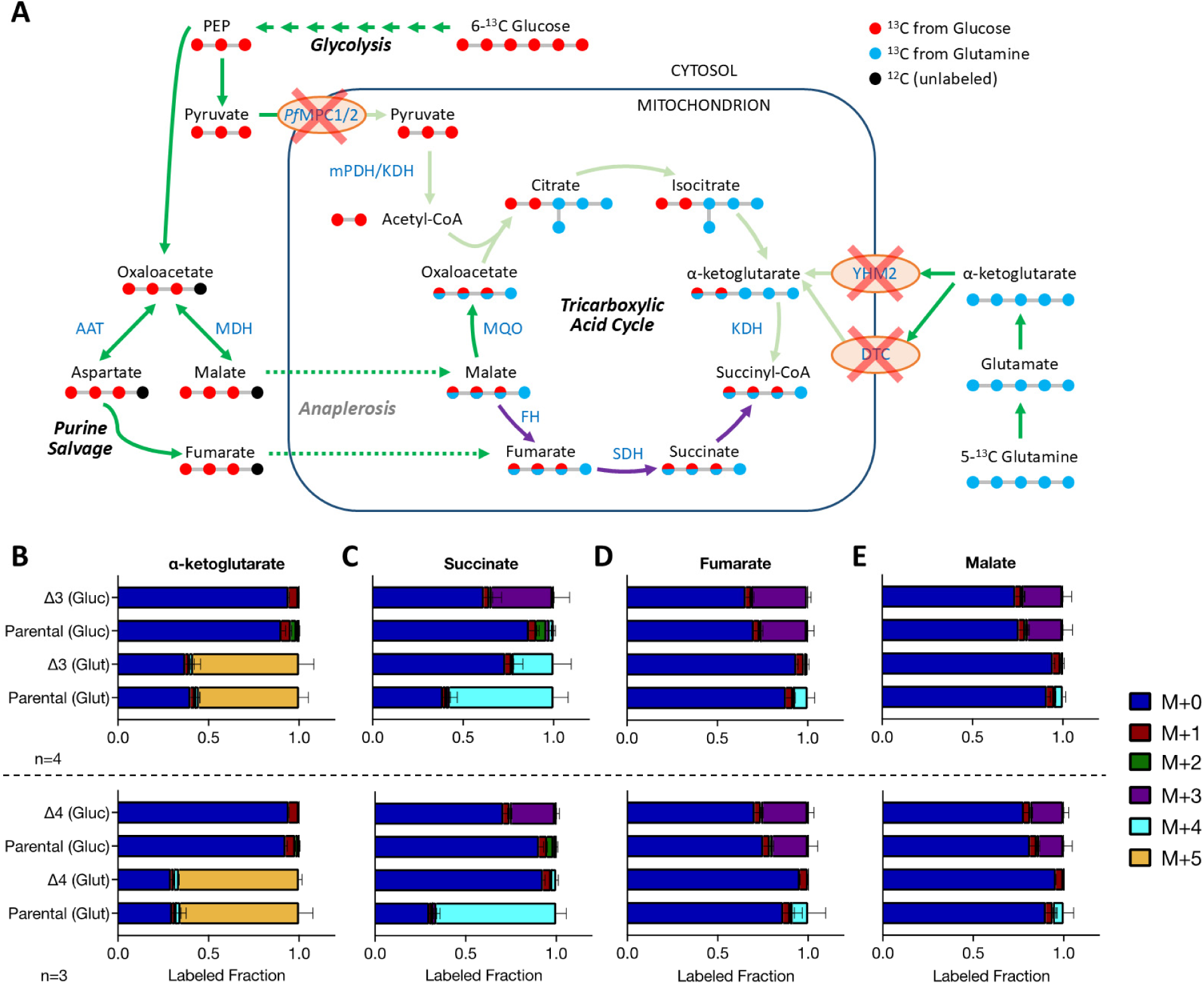
DTC and YHM2 control glutamine carbon incorporation into TCA cycle metabolites. (A) Schematic depicting the labeling pattern of mitochondrial metabolites in Δ4 parasites after short-term incubation with uniformly labeled ^13^C-glucose (solid red circles) or ^13^C-glutamine (solid blue circles). Black circles represent unlabeled atoms. Circles colored with both red and blue denote carbons that may be derived from either ^13^C_6_-glucose or ^13^C_5_-glutamine, respectively. Arrows represent enzymatic reactions and are colored based on the known flow (dark green), reduced flow (light green), predicted flow (dotted green) or reverse flow (purple) of carbon atoms. Abbreviations: PEP – phosphoenolpyruvate; AAT – aspartate aminotransferase; MDH – malate dehydrogenase; mPDH – mitochondrial pyruvate dehydrogenase; – KDH α-ketoglutarate dehydrogenase; SDH – succinate dehydrogenase; FH – fumarate dehydrogenase; MQO – malate-quinone oxidoreductase. (B-E) Fraction of isotopically labeled metabolites when parental, Δ3 or Δ4 parasites are incubated with labeled glucose (Gluc) or glutamine (Glut). Color coding denotes the mass (M) of the parent compound plus the mass shift from the incorporation of heavy carbon isotope (^13^C). Labeling data are expressed as the fraction of the total metabolite pool observed in ‘n’ independent experiments. Error bars represent the standard deviations from the mean.

Interestingly, succinate was found to contain three labeled carbons from glucose (M+3) in Δ3 and Δ4 parasites but not in the parental or Δ2 lines (**Fig. 4C**, **Fig. S4B**). This M+3 succinate is likely derived from an anaplerotic pathway in the cytosol that produces M+3 oxaloacetate from the glycolytic intermediate phosphoenolpyruvate (Cobbold et al., 2013; MacRae et al., 2013; Storm et al., 2014). Oxaloacetate can be converted into malate or aspartate in the cytoplasm, with the latter generating fumarate during purine salvage (**Fig. 4A**) (Bulusu et al., 2011; Storm et al., 2014). Anaplerotic glucose carbon may enter the TCA cycle as malate or fumarate. We speculate that the reduced level of α-ketoglutarate in the mitochondria of Δ3 and Δ4 parasites leads to a reversal in the direction of TCA cycle reactions downstream of KDH, with malate successively converted to fumarate and succinate by fumarate hydratase and succinate dehydrogenase, respectively. The end result is an accumulation of M+3 succinate in the Δ3 and Δ4 parasite lines (**Fig. 4A**).

### Glucose limitation does not exacerbate growth defect of parasites lacking multiple carboxylate transporters

Despite the observed perturbations in central carbon metabolism, we did not see a large fitness loss in the transporter mutants (**Fig. 1D**, **Fig. 3D-E**). We wondered if this was an artifact of culturing parasites in medium with roughly twice the glucose concentration (11 mM) compared to that found in human blood (4.0 – 6.5 mM). We therefore compared the growth of Δ4 and parental parasites in regular culture medium (11 mM glucose) and low glucose (3.67 mM glucose) medium over an 8-day period, with media replenishment every other day. While glucose levels did not alter growth kinetics during the first cycle, both parasite lines exhibited similarly reduced multiplication rates over the next three cycles in low glucose medium (**Fig. 5**). We therefore conclude that the high levels of glucose in regular culture medium are not masking fitness defects in Δ4 parasites.

**Fig. 5.**
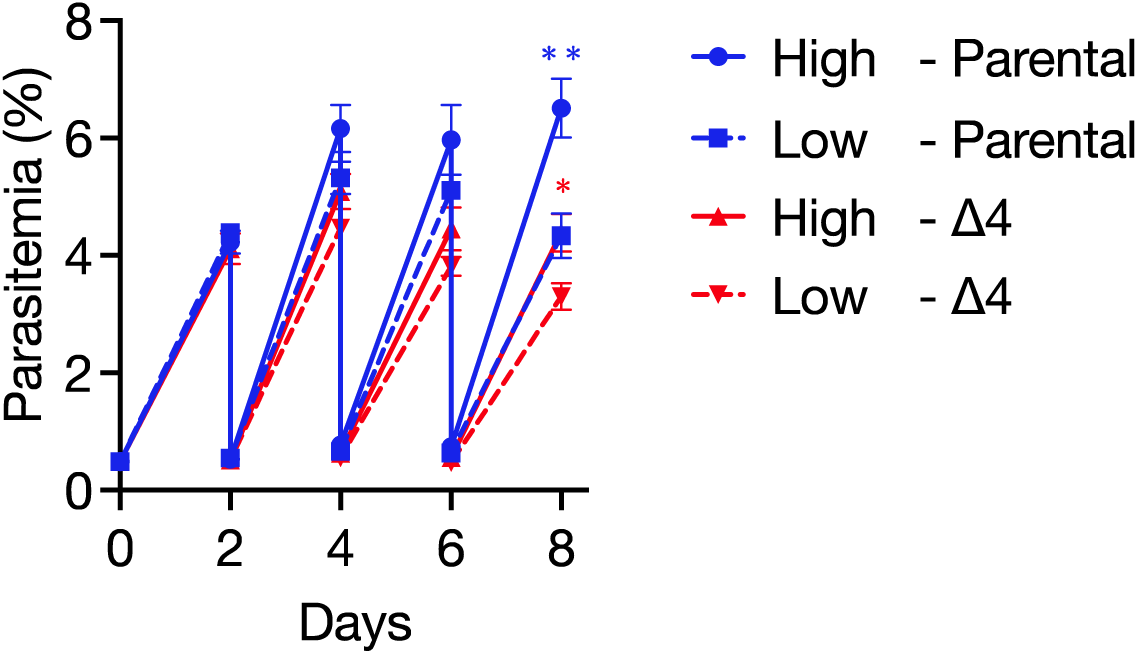
Carboxylate transporter-deficient parasites do not exhibit heightened sensitivity to low glucose conditions. Growth of parental and Δ4 parasites in low (3.67 mM) and high (11 mM) glucose media was monitored by flow cytometry over an 8-day period. Cultures were seeded at 0.5% parasitemia on day 0 and diluted 1:8 every other day. Two biological experiments were conducted in quadruplicate. Error bars represent the standard deviations from the mean. Statistics: Two-way ANOVA using Prism 10 [Graphpad Software, Inc], with Šídák’s multiple comparison test; *, p ≤ 0.05; **, p ≤ 0.01.

### Deletion of mitochondrial transporters does not alter parasite responsiveness to dihydroartemisinin

The mitochondrion of the malaria parasite is a key target of artemisinins. These drugs exert their antimalarial effects in part by inducing rapid mitochondrial membrane depolarization (Peatey et al., 2015; Wang et al., 2010). Arteminisin-resistant parasites have been found to exhibit altered mitochondrial energy metabolism, with a broad upregulation of genes linked to mitochondrial transport and pathways related to pyruvate, glutamate, and acetyl-CoA metabolism (Chen et al., 2014; Mok et al., 2021; Yu et al., 2023). In particular, transcription of the YHM2 transporter gene is upregulated in several clinically resistant isolates; however, whether this single-handedly alters responsiveness to artemisinin has not been established (Bonive-Boscan et al., 2023; Tripathi et al., 2023; Zhang et al., 2022; Zhu et al., 2022).

We utilized the ΔYHM2 and the dual copy parasite lines to determine the impact of increased gene dosage of YHM2 versus no expression on sensitivity to dihydroartemisinin (DHA). Employing standard 72-h proliferation assays, we found that the two transgenic strains and the NF54^attB^ parent had comparable IC_50_ values for DHA (Parental – 1.9 nM; YHM2-SFG – 1.8 nM; ΔΥΗΜ2 – 2.5 nM) (**Fig. 6A-B**). Since the Δ4 transporter mutant exhibits altered pyruvate and glutamate metabolism, we also assessed its sensitivity to DHA. We did not observe a notable shift in the IC_50_ value (Parental – 1.7 nM; Δ4 – 1.4 nM) (**Fig. 6C**).

**Fig. 6.**
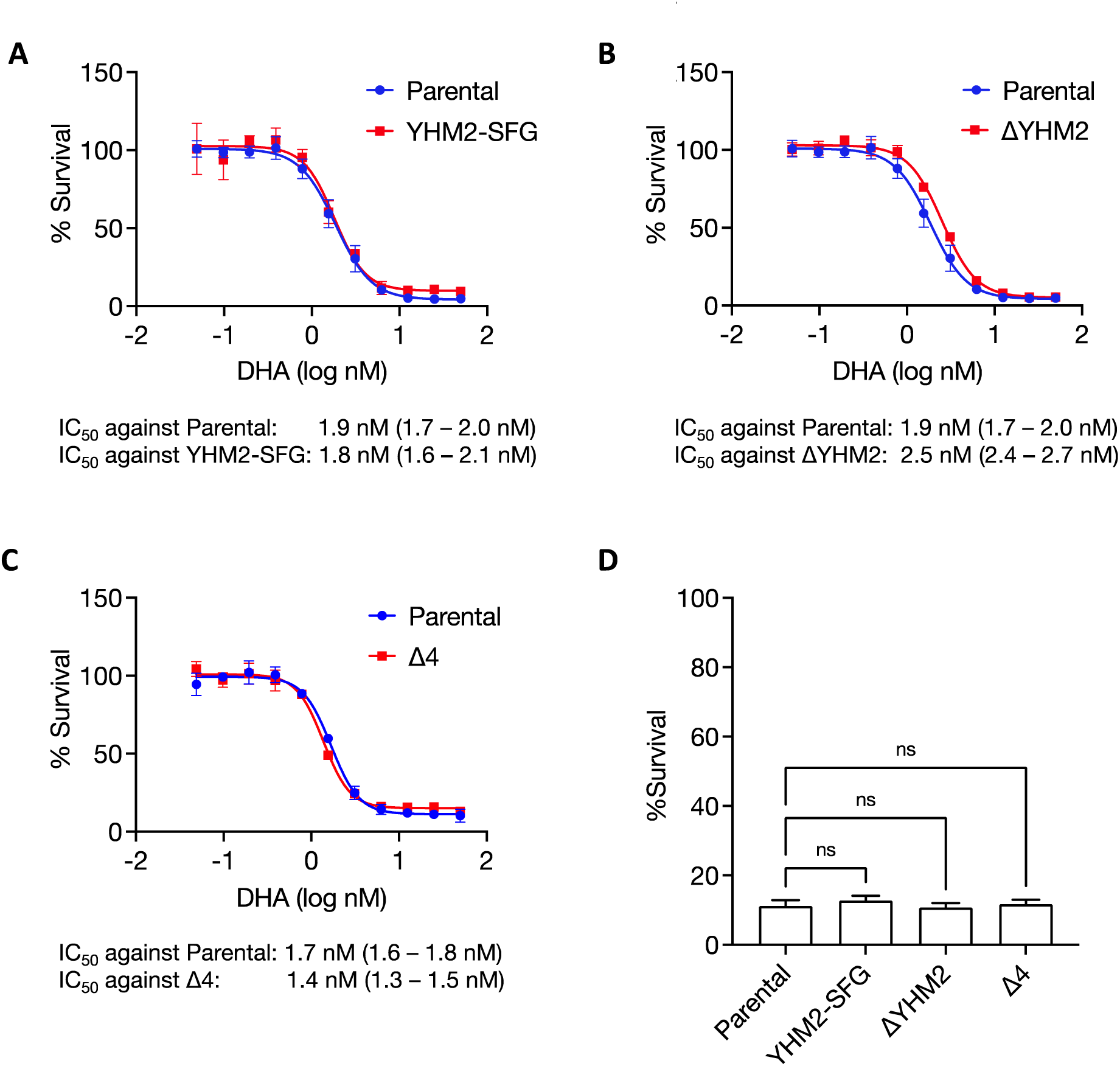
Mitochondrial transporter mutants do not show altered susceptibility to dihydroartemisinin (DHA). Parasites that (A) express a second copy of YHM2 under the constitutively active Cam/HOP promoter, (B) contain a YHM2 gene deletion, or (C) contain deletions in *Pf*MPC1, *Pf*MPC2, DTC and YHM2 genes, were exposed to various concentrations of dihydroartemisinin (DHA) for 72 h followed by measurement of parasitemia via flow cytometry. Parental parasites were employed as control. Two independent biological experiments with quadruplicate samples were conducted. Error bars represent the standard deviation from the mean. The parental data are identical in Fig. 6A and 6B since these experiments were conducted concurrently. The calculated IC_50_ values for DHA (with 95% confidence intervals in brackets) are listed below the graphs. C) Survival percentages from a ring-stage assay for parental, YHM2-SFG, ΔYHM2, and Δ4 parasites are depicted. Two independent biological experiments were conducted with quadruplicate samples and parasitemia was measured by flow cytometry at 66 h. Error bars represent the standard deviations from the mean. Statistics: IC_50_ values were obtained using a nonlinear variable slope four-parameter regression curve fitting model in Prism 10 [GraphPad Software Inc.]. One-way ANOVA with Dunnett’s correction in Prism 10 was used to compare differences between RSA survival percentages; ns – not significant.

Artemisinin resistance is characterized by delayed parasite clearance in patients and increased survival of early ring stage parasites *in vitro* (Amaratunga et al., 2012; Dondorp et al., 2009; Witkowski et al., 2010). The latter can be assessed through a ring-stage survival assay (RSA) in which 0-3 h rings are exposed to 650 nM DHA for 6 h and their recovery is measured after 66 h (Witkowski et al., 2013). We conducted parallel RSAs for the mutant and parent parasites and found that neither YHM2 overexpression nor deletion of one or multiple mitochondrial transporters influenced the ring-stage survival percentage (**Fig. 6D**). Therefore, our data suggest that changes in mitochondrial transporter expression alone do not alter the susceptibility of *P. falciparum* to DHA.

## DISCUSSION

Acetyl-CoA is a universally required metabolite that orchestrates numerous cellular processes mainly by donating its two-carbon acetyl group to recipient molecules (Shi and Tu, 2015). Because it is membrane-impermeable, distinct pools of acetyl-CoA are typically maintained by enzymes in various subcellular compartments. In the mitochondria of most eukaryotes, acetyl-CoA is derived from glucose, fatty acid, and branched-chain amino acid catabolism, and it undergoes complete oxidation through the TCA cycle to generate ATP and CO_2_. Cytosolic acetyl-CoA, which is required for lipid and isoprenoid synthesis, can be sourced in at least two ways: from the export and cleavage of mitochondrial citrate by ATP-citrate lyase, or from acetate by an acetyl-CoA synthetase (ACS). Acetyl-CoA can enter the nucleus through the nuclear pore or can be additionally generated by a resident ACS to regulate gene expression via acetylation of histones and transcription factors. The *P. falciparum* genome appears to encode some, but not all, of the enzymes responsible for acetyl-CoA production. During asexual blood stage development, the mitochondrion serves as the principal source of acetyl-CoA for the entire parasite (Nair et al., 2023). The pathways for BCAA degradation and fatty acid oxidation are either missing or incomplete, which leads us to surmise that glycolytic pyruvate is the primary substrate for acetyl-CoA synthesis in the mitochondrion (Cobbold et al., 2013; Gardner et al., 2002; MacRae et al., 2013; van Dooren et al., 2006). Indeed, inactivation of mPDH and KDH, the enzymes required for the conversion of pyruvate to acetyl-CoA, is lethal for parasites (Nair et al., 2023). Given the critical role of pyruvate in the mitochondrion, this study aimed to determine how pyruvate is mobilized from the cytoplasm into the organelle in malaria parasites.

We targeted two putative mitochondrial pyruvate carrier proteins – *Pf*MPC1 and *Pf*MPC2 –for genetic and metabolic studies in *P. falciparum*. While gene deletion of one or both proteins surprisingly did not strongly affect parasite viability in red blood cells, our isotopic labeling analyses revealed that the ^13^C label from glucose was considerably diminished in acetyl-CoA. However, this reduction was not as severe as seen previously in the mPDH/KDH double-deletion mutant, which suggested that one or more redundant mechanisms facilitate glucose carbon entry into the mitochondrion (Nair et al., 2023). To determine if other mitochondrial carriers can transport pyruvate, we turned our attention to two carboxylate carrier proteins, DTC and YHM2. Single and combinatorial deletion mutants were successfully obtained with modest fitness costs. Treatment of Δ3 (Δ*Pf*MPC1/*Pf*MPC2/DTC) and Δ4 (ΔMPC1/*Pf*MPC2/DTC/YHM2) mutants with ^13^C_6_-glucose demonstrated that the absence of DTC and YHM2 does not noticeably affect the levels of pyruvate-derived acetyl-CoA. Taken together, these results establish that *Pf*MPCs are the primary mitochondrial pyruvate carriers, but in their absence, an alternative path exists for some cytosolic pyruvate to enter the mitochondrion.

Although additional details of acetyl-CoA synthesis in the *P. falciparum* mitochondrion remain to be established, we were able to gain new insights into how another major carbon source, α-ketoglutarate, is imported into the organelle. Our ^13^C_5_-glutamine labeling data show that DTC plays a key role in facilitating glutamine carbon entry into the TCA cycle as α-ketoglutarate. In *P. falciparum*, glutamine is initially converted by glutamine synthetase to glutamate, which can then be converted to α-ketoglutarate in the cytosol by aspartate aminotransferase or by glutamate dehydrogenases (Storm et al., 2011). A putative GABA transaminase is speculated to convert glutamate to α-ketoglutarate in the mitochondrion, but our results indicate that the bulk of glutamine carbon reaches the organelle as α-ketoglutarate, not as glutamate (Cobbold and McConville, 2014). This is evidenced by the observation that DTC deletion did not impact glutamine incorporation into α-ketoglutarate, but it affected the labeling of other TCA cycle metabolites. Additional deletion of YHM2 exacerbated this metabolic phenotype, consistent with both transporters importing α-ketoglutarate and conversion of glutamine to α-ketoglutarate occurring prior to mitochondrial import.

The anaplerotic influx of glucose carbon into the TCA cycle has been attributed to DTC, which can transport oxaloacetate, malate and to a lesser extent, fumarate *in vitro* (Ke et al., 2015; Nozawa et al., 2011). Interestingly, DTC deletion did not block anaplerotic M+3 carbon assimilation into TCA cycle intermediates. On the contrary, we noted a large increase in M+3 succinate species when compared with parental parasites. We attribute this buildup to a reversal in the flow of glucose carbon from M+3 malate/fumarate in response to low levels of α-ketoglutarate in the mitochondrion. So, if DTC is not the relevant transporter, how does anaplerotic glucose carbon get into the mitochondrion, and in what form? A previous study indicated that M+3 oxaloacetate does not enter the organelle; the unknown mitochondrial carrier must therefore import anaplerotic fumarate or malate (Ke et al., 2015). If the carrier exhibits broad specificity for carboxylates, it might also serve as the redundant pyruvate transporter.

Although YHM2 is annotated as a citrate/α-ketoglutarate antiporter, it is difficult to determine whether the deletion of YHM2 impacts citrate levels in the mitochondrion and the cytoplasm, since our data reflect metabolite levels in the infected red blood cell as a whole. Even if YHM2 exports citrate from the mitochondrion, the fate of citrate in the cytoplasm is uncertain since malaria parasites do not appear to contain an ATP-citrate lyase to transform citrate into acetyl-CoA (Gardner et al., 2002). The yeast YHM2 homolog supports biosynthetic reactions and antioxidant defense by coupling citrate/α-ketoglutarate transport to NADPH generation (Castegna et al., 2010). While it is unclear if NADPH can be formed from citrate interconversion in the cytoplasm of *P. falciparum*, YHM2 has been recently linked to the artemisinin resistance phenotype, with such parasites demonstrating increased tolerance to drug-induced oxidative damage (Rocamora et al., 2018). The YHM2 gene is one of several genes associated with mitochondrial energy metabolism that are upregulated in resistant clinical isolates (Tripathi et al., 2023; Zhu et al., 2022). We found that deleting or expressing a second copy of YHM2 did not alter parasite susceptibility to DHA, indicating that YHM2 is not sufficient to significantly alter artemisinin resistance.

The metabolite data collected from our series of combinatorial deletion mutants allowed us to propose substrate specificities for several mitochondrial carriers. Our results also underscore the high degree of functional redundancy and plasticity in *P. falciparum* parasites, as evidenced by their ability to survive the loss of four carrier proteins. The gene deletion results presented in this study are not entirely concordant with a forward genetic screen for essentiality in *P. falciparum*, in which *Pf*MPC1, *Pf*MPC2 and DTC were all assigned low mutability scores (**Table 1**) (Zhang et al., 2018). However, the relatively small sizes of the coding sequences of these genes (*Pf*MPC1: 318 bp, *Pf*MPC2: 387 bp, DTC: 954 bp) decrease the predictive capability of the screen. On the other hand, YHM2 was identified as dispensable in both *P. falciparum* and *P. berghei* screens (Schwach et al., 2015; Zhang et al., 2018). Therefore, targeted gene deletion studies remain important to determine the functional role of genes expressed during the asexual blood stage.

**Table 1.** Essentiality of mitochondrial transporter genes from forward and reverse genetic approaches. a – Zhang et al., 2018; b – Schwach et al., 2015; c – Claros and Vincens, 1996; d – Esveld et al., 2021. * The *P. falciparum* genome-wide screen for essential genes (Zhang et al., 2018) utilized MFS (mutagenesis fitness score) and MIS (mutagenesis index score) ranging from 0 to 1 to predict gene mutability. Low MFS and MIS scores indicate gene essentiality. ** MitoProtII scores (ranging from 0 to 1) and PlasmoMitoCarta scores (ranging from -17.2747 to 12.486) predict mitochondrial localization of *Plasmodium* proteins. Proteins with high MitoProtII scores and high PlasmoMitoCarta scores have a higher probability of localizing to the mitochondrion.

| Gene ID<br>(PF3D7_) | Gene name | Essentiality in asexual blood-stage *Plasmodium* |  |  |  | Mitochondrial Localization |  |
| --- | --- | --- | --- | --- | --- | --- | --- |
|  |  | *P. falciparum* (a) |  | *P. berghei* (b) | *P. falciparum* (this study) | MitoProtII (c)** | PlasmoMitoCarta (d)** |
|  |  | MFS* | MIS* |  |  |  |  |
| 1340800 | Mitochondrial pyruvate carrier 1 (*PfMPC1*) | -3.235 | 0.119 | Not studied | Dispensable | 0.8878 | 5.97197 |
| 1470400 | Mitochondrial pyruvate carrier 2 (*PfMPC2*) | -2.167 | 0.331 | Not studied | Dispensable | 0.6315 | 6.87333 |
| 0823900 | Di/tricarboxylic acid carrier (DTC) | -2.799 | 0.162 | Not studied | Dispensable | 0.2626 | 3.54392 |
| 1223800 | Citrate/α-ketoglutarate transporter (YHM2) | -0.31 | 1 | Dispensable | Dispensable | 0.1695 | 1.31981 |

In summary, our study demonstrated that pyruvate is imported into the mitochondrion of *P. falciparum* parasites via *Pf*MPC1 and *Pf*MPC2 and at least one additional path. We also found that α-ketoglutarate is transported into the mitochondrion by DTC and YHM2 to feed the TCA cycle. It remains to be determined how acetyl-CoA is exported from the mitochondrion for utilization in other intracellular compartments. The importance of acetyl-CoA metabolism in malaria parasites warrants further experimentation to identify and characterize novel mitochondrial carriers that support the synthesis and utilization of this critical metabolite.

## METHODS

### Parasite culture

*P. falciparum* NF54^attB^ parasites were cultured in red blood cells (RBCs) at 2% hematocrit in complete medium with AlbuMAX II (CMA). CMA is prepared from RPMI-1640 medium with L-glutamine (USBiological Life Sciences, R8999) and is supplemented with 20 mM HEPES, 0.2% sodium bicarbonate, 12.5 μg/mL hypoxanthine, 5 g/L AlbuMAX II (ThermoFisher Scientific, 11021037) and 25 μg/mL gentamicin. Parasite cultures were maintained at 37°C in a gas mixture containing 94% N_2_, 3% O_2_ and 3% CO_2_.

### Plasmid construction

Localization constructs were derived from the previously published p15-EcDPCK-mCherry plasmid (Swift et al., 2021). The mCherry tag was removed by digesting the plasmid with BsiWI and AflII, and replaced with an SFG sequence that was amplified using primers listed in **Table S1** and the p15-Mev-aSFG plasmid as template (Swift et al., 2020b). The SFG amplicon was digested with BsiWI and AflII and ligated with the cut p15-EcDPCK plasmid using T4 DNA Ligase (NEB, M0202). The EcDPCK gene was replaced with *Pf*MPC1, *Pf*MPC2, YHM2 or DTC coding sequences by digesting the plasmid with AvrII and BsiWI. The coding sequences were amplified from *P. falciparum* NF54^attB^ genomic DNA (mRNA for *Pf*MPC1 and *Pf*MPC2 to exclude introns) with primers included in **Table S1**. The amplicons were digested with BsiWI and AflII and ligated into the digested p15-SFG plasmid.

Gene deletion constructs for *Pf*MPC1, *Pf*MPC2, DTC and YHM2 were generated using several versions of pRSng repair plasmids with different drug resistance cassettes (Rajaram et al., 2022; Swift et al., 2020b). Homology arms 1 (HA1) and 2 (HA2) for the genes were amplified using primers listed in **Table S1**. The HA1 sequence was inserted into the NotI site and HA2 into the NgoMIV site of the repair plasmid. For individual knockouts of the *Pf*MPC genes, we used the original pRSng plasmid which contains a hDHFR (human dihydrofolate reductase) resistance cassette that confers resistance to the drug WR99210 (Swift et al., 2020b). For deletion of *Pf*MPC1 in the Δ*Pf*MPC2 background, hDHFR was replaced with blasticidin deaminase (BSD) to generate a *Pf*MPC1-pRSngBD plasmid. Homology arms for DTC were cloned into a pRSng-Neo plasmid containing a neomycin cassette, providing resistance to G418 (Rajaram et al., 2022). Finally, homology arms for YHM2 were cloned into a pRSng-Mev plasmid which contains a mevalonate cassette that provides resistance to fosmidomycin. To create the pRSng-Mev plasmid, the pRSng and p15-Mev-aSFG plasmids were digested with BamHI and HindIII to replace the hDHFR cassette in pRSng with the Mev-aSFG cassette from p15-Mev-aSFG using ligation (Swift et al., 2020b).

To generate plasmids with guide RNA sequences that target *Pf*MPC1, *Pf*MPC2, DTC and YHM2 for gene deletion, the pCasG-LacZ vector was digested with the type IIS restriction enzyme BsaI (Rajaram et al., 2020). Guide RNAs were synthesized as oligonucleotides (**Table S1**), annealed, and inserted into the cut plasmid. Unless specified, plasmid construction relied on ligation-independent cloning with In-Fusion (Takara, 638944). Restriction enzymes were sourced from New England Biolabs.

### Parasite transfections

To determine the subcellular localization of *P. falciparum* transporter proteins, 350 μL volumes of RBCs were electroporated using the GenePulser XCell system (Bio-Rad) with: 50 μg of the pINT plasmid (Nkrumah et al., 2006) that expresses the Bxb1 integrase, and 65 μg of the p15-SFG plasmid expressing either *Pf*MPC1, *Pf*MPC2, DTC or YHM2 fused to a C-terminal SFG tag. The electroporated RBCs were mixed with 1 mL of NF54^attB^ parasite culture at 2-3% parasitemia and maintained in 10 mL of CMA for 2 days. They were then transferred to selective media containing 2.5 nM WR99210 for 7 days. The cultures were then returned to CMA until blood smears turned parasite-positive, whereupon WR99210 was added back to the medium.

To generate deletion strains, 350 μL volumes of RBCs were electroporated with 65 μg each of the repair plasmid and the appropriate pCasG-gRNA plasmid. The electroporated RBCs were mixed with 1 mL of parasite culture at 2-3% parasitemia and maintained in 10 mL of CMA for 2 days. They were then transferred to selective media containing 1.5 μM DSM1 and either 2.5 nM WR99210 (for individual deletions of *Pf*MPC1 and *Pf*MPC2), 0.5 mg/mL G418 (for DTC knockout), or 25 μM fosmidomycin plus 50 μM mevalonolactone (for YHM2 knockout) for 7 days. Cultures were then switched to CMA until parasites reappeared, after which they were transferred to CMA containing WR99210 (Δ*Pf*MPC1 and Δ*Pf*MPC2) or G418 (ΔDTC). To delete *Pf*MPC1 in the Δ*Pf*MPC2 line, RBCs were electroporated with the pRSngBD-*Pf*MPC1 plasmid and the same pCasG-gRNA plasmid employed earlier. Two days post transfection, the culture was selected with 2.5 μg/mL blasticidin for 7 days. Culture media then was replaced with CMA, and maintained as such until parasite reappearance, at which point blasticidin was reintroduced into the media.

All transgenic parasite lines were cloned by limiting dilution. Drugs were acquired from the following vendors: DSM1 – Calbiochem (CAS 92872-51-0); WR99210 – Jacobus Pharmaceutical Company, Inc.; Blasticidin – Corning (30-100-RB); G418 – Corning (61-234-RG), Fosmidomycin – ThermoFisher Scientific (FR-31564), Mevalonolactone – Sigma-Aldrich (M4667).

### Genotyping PCR

Three μL volumes of parasite culture were used in 25 μL PCR reactions with Phusion High-Fidelity DNA polymerase (NEB, M0530). Primer pairs and anticipated sizes of PCR products are indicated in **Fig. S1** and **Fig. S2**. Primer sequences can be found in **Table S1**.

### Growth assays

The growth rates of gene knockout lines were compared against the parental line to determine if gene deletions imparted fitness costs. Parasites were used to seed 200 μL culture volumes/well in a 96-well plate (ThermoFisher Scientific, 267427) at a 0.5% starting parasitemia and 2% hematocrit. The plates were incubated in chambers gassed with 94% N_2_, 3% O_2_, 3% CO_2_ at 37°C. After 48 h (the duration of an asexual intraerythrocytic developmental cycle), final parasitemia was determined by flow cytometry. This process was repeated over at least four growth cycles, with an eight-fold dilution of cultures every 48 h to prevent overgrowth. Samples were prepared for flow cytometry as described previously (Swift et al., 2020b). Briefly, parasites were diluted 1:5 in PBS and stored in a 96-well plate at 4 °C until analysis by flow cytometry. Parasites were stained by transferring 10 μL of the 1:5 dilutions to a 96-well plate containing 100 μL of 1x SYBR Green (Invitrogen, S7563) in PBS, and incubated for 30 minutes in the dark while shaking. Post-incubation, 150 μL of PBS was added to each well to dilute unbound SYBR Green dye. Stained cells were analyzed with an Attune NxT Flow Cytometer (ThermoFisher Scientific) at a running speed of 25 μL/minute to acquire 10,000 events. Experiments were conducted with quadruplicate samples.

### *P. falciparum* purification, isotopic labeling, and metabolomics analysis

Heparin-synchronized NF54^attB^ parental (Adjalley et al., 2011) and associated knockout strains of *P. falciparum* parasites were cultured in standard conditions. At approximately 24-30 hours post invasion, healthy trophozoite stage parasites were purified via magnet activated cell sorting using a SuperMACS™ II Separator (Miltenyi Biotec, 130-044-104) and a D column (Miltenyi Biotec, 130-041-201). The samples of ∼95-100% pure trophozoite-infected erythrocytes were plated at 0.2% hematocrit in standard culture media and incubated for a one-hour recovery period. After recovery, the parasites were pelleted, the standard culture media supernatant was discarded, and parasites were resuspended in complete media made with either glucose-dropout (Gibco, 11879-020) or glutamine-dropout (Gibco, 21870-076) base RPMI-1640 supplemented with 25mM HEPES pH 7.4 (Millipore Sigma, H4034), 2 g/L sodium bicarbonate (Fisher Scientific, BP328), 10 mg/L hypoxanthine (Fisher Scientific, AC122010250), 2.5 g/L AlbuMAX II (Thermo Fisher Scientific, 11021029), 50 mg/L gentamicin sulfate (Bio Basic, GB0217). 11 mM ^13^C_6_-glucose (Cambridge Isotope Laboratories, CLM-1396-1) or 2 mM ^13^C_5_-glutamine (Cambridge Isotope Laboratories, CLM-1822-H) was added to respective dropout media. Parasites were cultured in 6-well culture plates at 1×10^8^ cells/5 mL media in technical triplicate for a 2.5 hour labeling period.

After pelleting and washing in ice cold PBS pH 7.4, the infected erythrocytes were then resuspended in 1 mL of 90% HPLC-grade methanol with 0.5 μM [^13^C_4_,^15^N_1_] aspartate (Millipore Sigma, 607835) as an internal control. Supernatants were dried under nitrogen gas flow and stored at -70°C before being resuspended to 10^6^ former cells/µL in HPLC-grade water with 1µM chlorpropamide (Alfa Aesar, J64110) as an additional internal control for reverse phase UHPLC-MS analysis on a Thermo Exactive Plus Orbitrap. For each sample, 10 µL were injected through a XSelect HSS T3 2.5 µM C18 column (Waters, 186006151) and eluted using a 25 min gradient of 3% aqueous methanol, 15 mM acetic acid (Millipore Sigma, A6283), 10 mM tributylamine (Millipore Sigma, 90781), 2.5 µM medronic acid ion pairing agent (Millipore Sigma, M9508) (A) and 100% HPLC-grade methanol (B). Negative-ion mode, using a scan range of 85 to 1,000 *m/z* and a resolution of 140,000 at *m/z* 200 was used for ion detection.

Resulting .raw files were converted to .mzML format and centroided using the MSConvert software of the ProteoWizard package (Chambers et al., 2012). The .mzML files were then aligned, peaks picked, and peaks annotated (including isotopic variants) with El-MAVEN (Agrawal et al., 2019). The chlorpropamide internal standard intensity as well as pooled quality control (QC) samples were used to evaluate technical reproducibility of ion detection across the run. EL-MAVEN-called peaks were then manually curated based on the proximity to the standard retention time in the targeted library, signal/blank ratio, and peak shape. Integrated areas of acceptable peaks were exported from EL-MAVEN in a.csv file and further processing was performed using Microsoft Excel.

Chlorpropamide standard abundance was used to correct the peak areas per metabolite for instrument variation in ion detection during the run. Subsequently, the average blank signals were subtracted (or substituted to avoid negative abundances), and metabolites were filtered based on the relative standard deviation (RSD, Standard Deviation/Mean) of QC samples run regularly throughout data collection and the metabolites were excluded from analysis if RSD > 30% (**Supplemental File 2**). Raw data for all metabolites are available at the NIH workbench under submission Project ID: PR002196.

### Whole genome sequencing of clonal parasite lines

Parasites used for genomic DNA (gDNA) extraction were grown to 8 - 10 % schizonts in a 50-mL culture at 2% hematocrit. Cells were isolated by centrifugation (400 g for 3 minutes) before aspirating the supernatant and adding 50 mL of 0.1% saponin in PBS. Following 10 minutes of lysis, parasites were isolated by centrifugation (1,500 g for 10 minutes) and washed twice with 50 mL of PBS. The gDNA was extracted using a Qiagen DNeasy Blood and Tissue kit (69504) according to the provided protocol. Extracted gDNA was quantified using a Nanodrop spectrophotometer or a Qubit 2.0 fluorometer. Sequencing libraries for Δ2 and Δ3 lines were prepared with 1 μg of gDNA per sample using the TruSeq® DNA PCR-Free Low-Throughput Library Prep Kit (Illumina, 20015962) according to the manufacturer’s directions. The Δ2 and Δ3 libraries were sequenced on a HiSeq2500 Illumina platform. The sequencing library for the Δ4 line was prepared with 100 ng of gDNA using the Celero PCR Workflow and an Enzymatic Fragmentation DNA Seq Kit (Tecan, 30238472). This library was sequenced on the iSeq 100 Illumina platform. Genomic sequence data for the Δ2, Δ3 and Δ4 transgenic lines are available at the NCBI Sequence Read Archive under submission BioProject PRJNA1181605.

### IC_50_ and ring-stage survival assays for DHA

To perform dose-response experiments with DHA, late-stage parasites from the parental NF54^attB^, ΔYHM2, YHM2-SFG and Δ4 lines were magnetically sorted from mixed-stage cultures using MACS LS columns (Miltenyi Biotech, 130-042-401). The purified parasites were diluted to 1% parasitemia in 1.5% hematocrit and incubated with serial dilutions of DHA resuspended in DMSO to achieve a final DMSO concentration of 0.1%. After 72 h of incubation in standard culturing conditions, parasitemia was measured by flow cytometry as described above.

For ring-stage survival assays, cultures were synchronized by magnetic column enrichment of late-stage parasites over two consecutive ABS cycles. After a 3 h period for reinvasion, parasites were treated with 5% sorbitol before initiation of the assay to remove any late-stage parasites. The highly synchronized ring-stage parasites from NF54^attB^, ΔYHM2, YHM2-SFG and Δ4 lines were maintained for 6 hours in CMA containing 650 nM of DHA in 0.1% DMSO. Simultaneously, parasites were incubated in CMA with 0.1% DMSO only to serve as controls. Following the 6 h incubation, the cultures were washed three times in CMA to remove drug or DMSO. The washed parasites were further cultured for 66 h before measurement of parasitemia by flow cytometry. Survival rates for each parasite line were determined by calculating the percentage of surviving DHA-treated parasites to DMSO-treated parasites.

## ACKNOWLEDGEMENTS

This work was supported by NIH R01 AI168183 (awarded to S.T.P. and M.L.), the Johns Hopkins Malaria Research Institute, and the Bloomberg Family Foundation. K.R. was supported by the Johns Hopkins Discovery Award, G.R. was supported by a Penn State Eberly Research Fellowship, and J.T.M. was supported by training grant T32 DK120509. M.L. was also supported by the Eberly College of Science and the Huck Institutes of the Life Sciences at The Pennsylvania State University. The authors would like to acknowledge the Huck Institutes’ Metabolomics Core Facility (RRID:SCR_023864) for maintenance of the Thermo Exactive Plus.

