## Supplemental File 1 for "MULTIPLE, REDUNDANT CARBOXYLIC ACID TRANSPORTERS SUPPORT MITOCHONDRIAL METABOLISM IN *PLASMODIUM FALCIPARUM*"

**A**

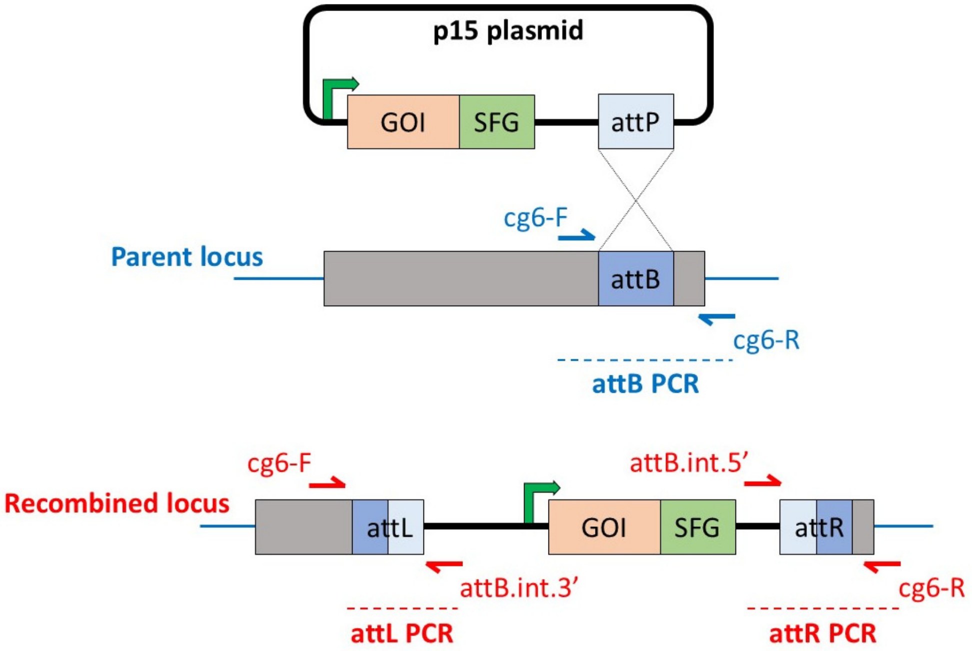


**B**

| Lane | PCR reaction | Primer Combination | | Product size (bp) | Lines |
|---|---|---|---|---|---|
| 1 | attL | cg6-F | attB.int.3’ | 334 | Recombinant |
| 2 | attR | attB.int.5’ | cg6-R | 1067 | |
| 3 | attB | cg6-F | cg6-R | – | |
| 4 | attL | cg6-F | attB.int.3’ | – | Parent |
| 5 | attR | attB.int.5’ | cg6-R | – | |
| 6 | attB | cg6-F | cg6-R | 1288 | |

**C**

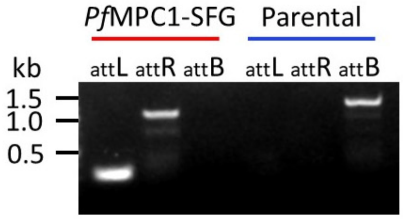


**D**

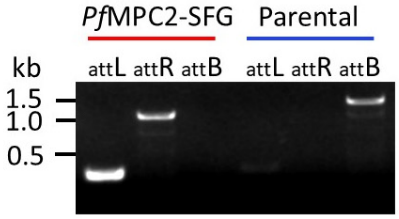


**Fig. S1**. **Generation of transgenic P.** ***falciparum*** **lines with SFG-tagged proteins of interest.** (A) The schematic depicts recombination of the p15-SFG plasmid with the NF54$^{attB}$ parasite genome. The p15-SFG plasmid is inserted into the cg6 locus of *P. falciparum* NF54$^{attB}$ by Bxb1 integrase-mediated recombination at the plasmid attP and genome attB sites. The modified locus contains the entire plasmid flanked by new attL and attR sites that are generated by recombination. Red and blue arrows indicate the positions of primers used for diagnostic PCR. The green arrow indicates the Cam/HOP promoter that drives expression of the transgene. Abbreviations: GOI – Gene Of Interest; SFG – Superfolder GFP. (B) Primer pairs for verifying integration of the p15-SFG plasmid into the cg6 locus are listed along with the expected sizes of the PCR products. Integration of (C) p15.*Pf*MPC1-SFG and (D) p15.*Pf*MPC2-SFG in NF54$^{attB}$ (parental) parasites was confirmed by PCR amplification (attL and attR products). The intact cg6 locus (attB product) was detected only in the parental (blue) and not in the transgenic lines (red).

**A**

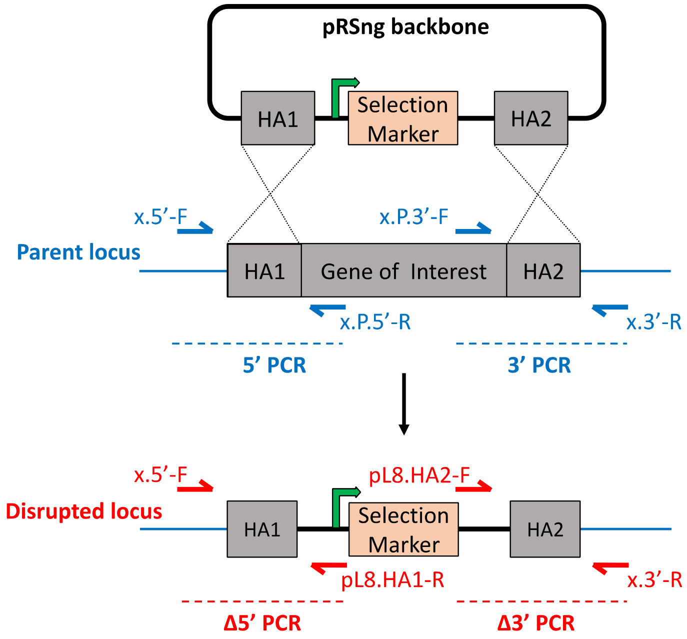


**B**

| Lane | PCR reaction | Primer Combination | | Δ*Pf*MPC1 product size (bp) | Δ*Pf*MPC2 product size (bp) | Lines |
|---|---|---|---|---|---|---|
| 1 | Δ5' | x.5'-F | pL8.HA1-R | 864 | 963 | KO |
| 2 | Δ3' | pL8.HA2-F | x.3'-R | 451 | 616 | |
| 3 | 5' | x.5'-F | x.P.5'-R | – | – | |
| 4 | 3' | x.P.3'-F | x.3'-R | – | – | |
| 5 | Δ5' | x.5'-F | pL8.HA1-R | – | – | Parent (P) |
| 6 | Δ3' | pL8.HA2-F | x.3'-R | – | – | |
| 7 | 5' | x.5'-F | x.P.5'-R | 820 | 1046 | |
| 8 | 3' | x.P.3'-F | x.3'-R | 581 | 949 | |

**Fig. S2. Schematic of CRISPR/Cas9-mediated disruption of mitochondrial transporter genes.** (A) The pRSng plasmid contains a selection marker flanked by sequences with homology to the gene of interest, indicated by dotted black lines. The green arrow indicates the location and directionality of the Cam/HOP promoter. Red and blue arrows indicate the positions of primers used for diagnostic PCR. The notation 'x' represents the targeted gene. Abbreviations: HA1 – Homology Arm 1; HA2 – Homology Arm 2; P – Parent. (B) Primer pairs for verifying *Pf*MPC1 and *Pf*MPC2 gene knockouts are listed along with the expected sizes of the PCR products.

**A**

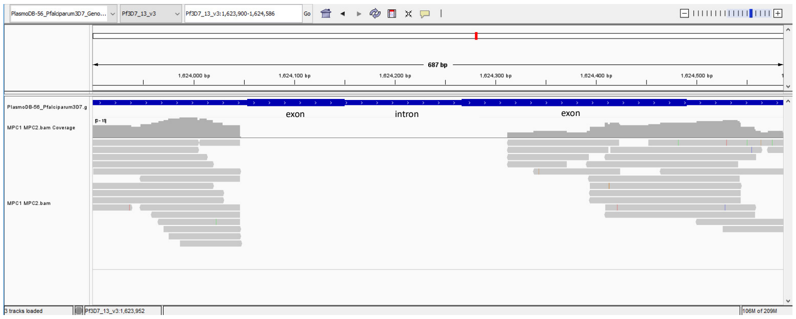


***Pf*MPC1** **PF3D7_1340800**
*Pf*MPC1 ORF: 1624054 – 1624490
HA1: 1623863 – 1624136
HA2: 1624313 – 1624549
Expected Δ: 1624137 – 1624312 (176 bp)
Observed Δ: 1624054 – 1624312 (257 bp)

**B**

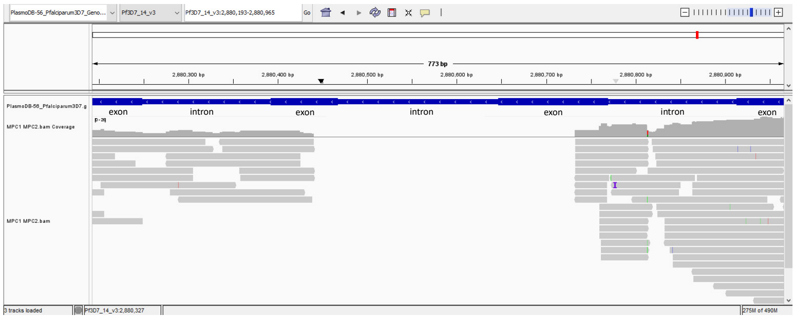


***Pf*MPC2** **PF3D7_1470400**
*Pf*MPC2 ORF: 2881012 – 2880157
HA1: 2881012 – 2880733
HA2: 2880440 – 2880198
Deleted region: 2880732 – 2880441 (292 bp)

**C**

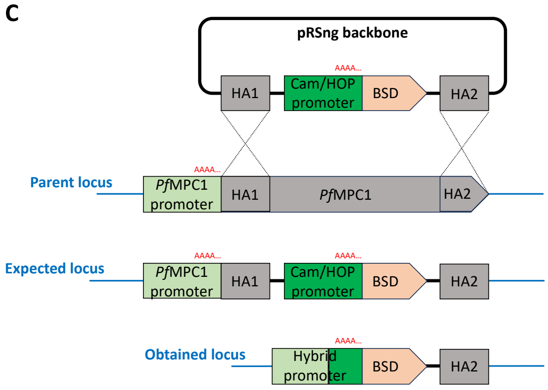


**Fig. S3**. **Confirmation of the Δ2 mutant genotype by whole genome sequencing.** Sequencing reads from the Δ2 genome were mapped to the *P. falciparum* reference genome using Bowtie2 (VEuPathDB Galaxy) (Alvarez-Jarreta et al., 2024) and visualized using Integrative Genomics Viewer (IGV) (Robinson et al., 2011). The images captured from IGV confirm the absence of reads within the open reading frames (ORF) of (A) *Pf*MPC1 in chromosome 13, and (B) *Pf*MPC2 in chromosome 14. The nucleotide positions of the *Pf*MPC1 and *Pf*MPC2 genes in their respective chromosomes, along with CRISPR homology arms HA1 and HA2 and the deleted regions in Δ2 parasites, are listed. (C) Schematic depicting the expected and obtained transgenic *Pf*MPC1 locus in Δ2 parasites. Whole genome sequencing revealed that the disrupted *Pf*MPC1 locus in Δ2 parasites differed from the anticipated result. A poly(A) sequence in the Cam/HOP promoter of the repair plasmid crossed over with a poly(A) site immediately upstream of the *Pf*MPC1 start codon. This resulted in the deletion of a large portion of the Cam/HOP promoter that drives the expression of a blasticidin deaminase (BSD) resistance cassette, forming a hybrid *Pf*MPC1-Cam/HOP promoter. The new promoter was still functional, allowing us to select for transgenic parasites using blasticidin. Despite this unexpected recombination event, the desired functional deletion of the *Pf*MPC1 gene was achieved.

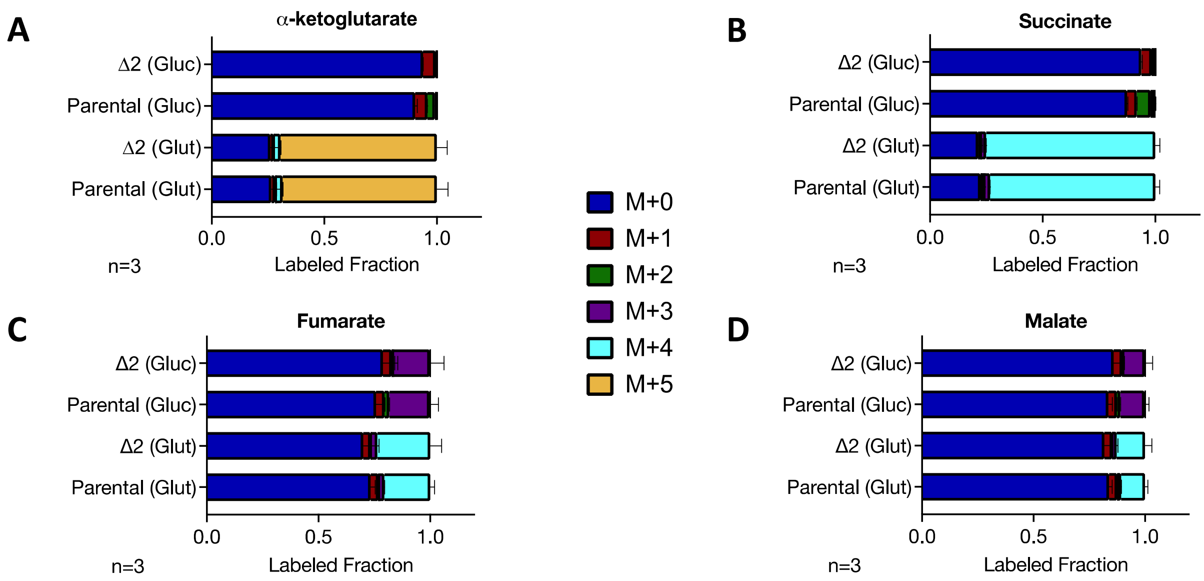


**Fig. S4**. ***Pf*MPC deletion does not heavily impact the production of TCA cycle metabolites from glucose.** Fraction of isotopically labeled metabolites when parental or Δ2 parasites are incubated with labeled glucose (Gluc) or glutamine (Glut). Color coding denotes the mass (M) of the parent compound plus the mass shift from the incorporation of heavy carbon isotope ($^{13}C$). Labeling data are expressed as the fraction of the total metabolite pool observed in 'n' independent experiments. Error bars represent the standard deviations from the mean.

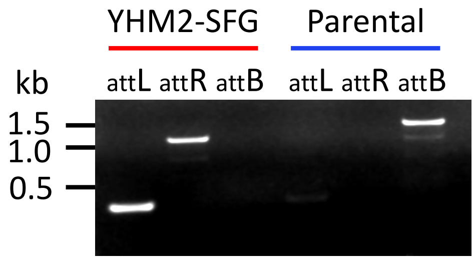


**Fig. S5**. **Generation of a transgenic P*. falciparum* line expressing SFG-tagged YHM2.** Integration of p15.YHM2-SFG in NF54$^{attB}$ (parental) parasites was confirmed by PCR amplification (attL and attR products). The intact cg6 locus (attB product) was detected only in the parental (blue) and not in the transgenic line (red).

| Lane | PCR reaction | Primer Combination | | ΔDTC product size (bp) | ΔYHM2 product size (bp) | Lines |
|---|---|---|---|---|---|---|
| 1 | Δ5’ | x.5’-F | pL8.HA1-R | 762 | 561 | KO |
| 2 | Δ3’ | pL8.HA2-F | x.3’-R | 1189 | 546 | |
| 3 | 5’ | x.5’-F | x.P.5’-R | – | – | |
| 4 | 3’ | x.P.3’-F | x.3’-R | – | – | |
| 5 | Δ5’ | x.5’-F | pL8.HA1-R | – | – | Parent (P) |
| 6 | Δ3’ | pL8.HA2-F | x.3’-R | – | – | |
| 7 | 5’ | x.5’-F | x.P.5’-R | 681 | 515 | |
| 8 | 3’ | x.P.3’-F | x.3’-R | 1293 | 672 | |

**Fig. S6. Confirmation of ΔDTC and ΔYHM2 mutations by PCR.** Primer pairs for verifying DTC and YHM2 gene deletions are listed along with the expected sizes of the PCR products. Gel images of the corresponding PCR products are shown in Fig. 3B-C.

A

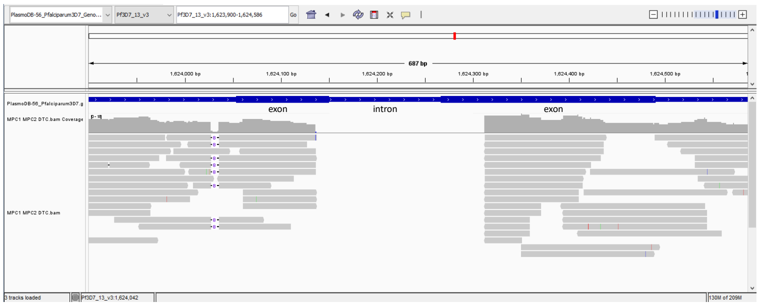


D

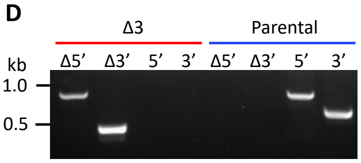


| | |
|---|---|
| ***Pf*MPC1** | **PF3D7_1340800** |
| *Pf*MPC1 ORF: | 1624054 – 1624490 |
| HA1: | 1623863 – 1624136 |
| HA2: | 1624313 – 1624549 |
| Expected Δ: | 1624137 – 1624312 (176 bp) |

B

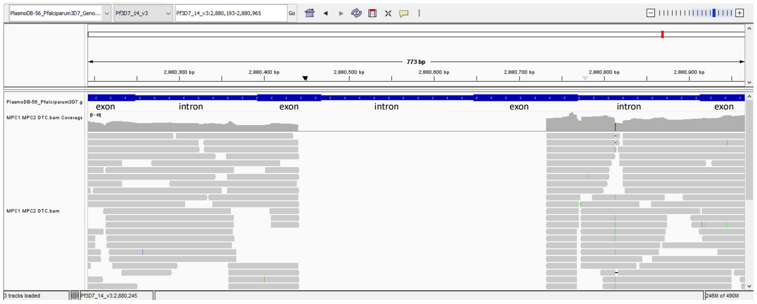


E

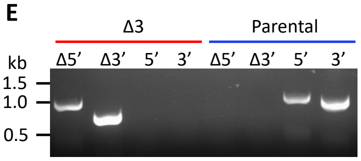


| | |
|---|---|
| ***Pf*MPC2** | **PF3D7_1470400** |
| *Pf*MPC2 ORF: | 2881012 – 2880157 |
| HA1: | 2881012 – 2880733 |
| HA2: | 2880440 – 2880198 |
| Deleted region: | 2880732 – 2880441 (292 bp) |

C

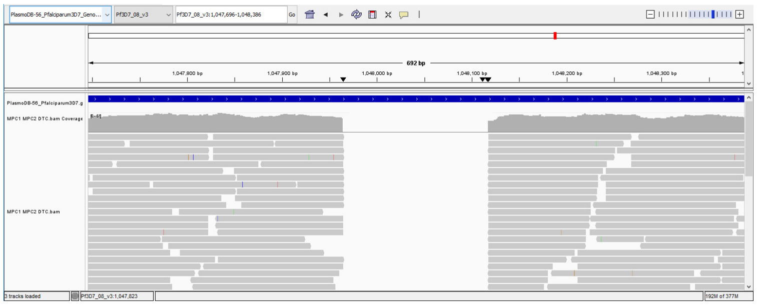


F

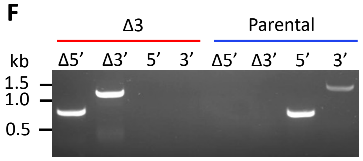


| | |
|---|---|
| **DTC** | **PF3D7_0823900** |
| DTC ORF: | 1047551 – 1048507 |
| HA1: | 1047550 – 1047964 |
| HA2: | 1048118 – 1048527 |
| Deleted region: | 1047965 – 1048117 (153 bp) |

**Fig. S7**. **Confirmation of the Δ3 mutant genotype by PCR and whole genome sequencing.** Sequencing reads from the Δ3 genome were mapped to the *P. falciparum* reference genome using Bowtie2 (VEuPathDB Galaxy) (Alvarez-Jarreta et al., 2024) and visualized using Integrative Genomics Viewer (IGV) (Robinson et al., 2011). The images captured from IGV confirm the absence of reads within the open reading frames (ORF) of (A) *Pf*MPC1 in chromosome 13, (B) *Pf*MPC2 in chromosome 14, and (C) DTC in chromosome 8. Gene deletion of (D) *Pf*MPC1, (E) *Pf*MPC2, and (F) DTC in the Δ3 mutant was confirmed by PCR amplification (Δ5' and Δ3' products). Intact gene loci (5' and 3' products) were detected only in the $NF54^{attB}$ parental line (blue) and not in the Δ3 line (red). The nucleotide positions of the *Pf*MPC1, *Pf*MPC2 and DTC genes in their respective chromosomes, along with CRISPR homology arms HA1 and HA2 and the deleted regions in Δ3 parasites, are listed.

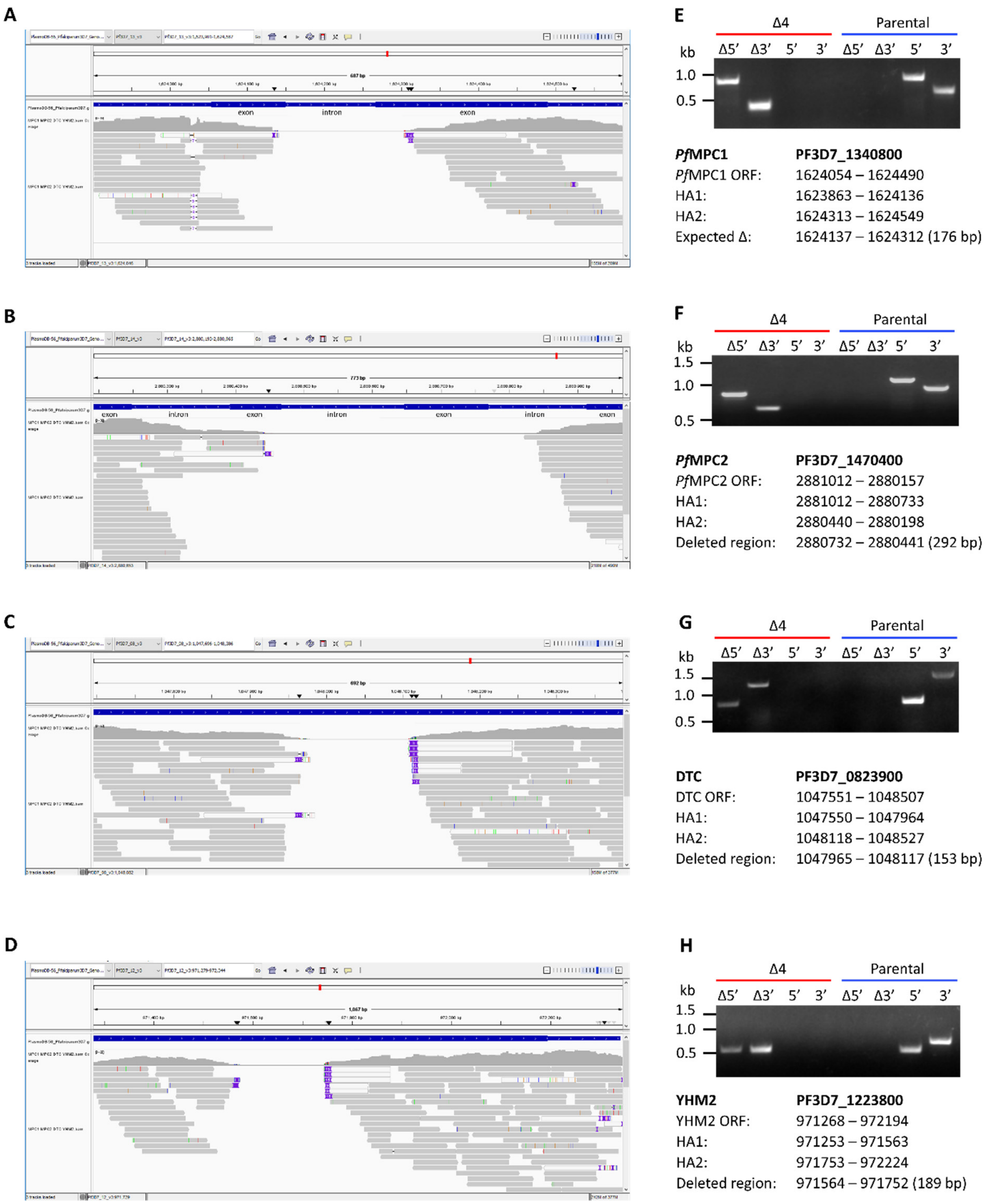


**Fig. S8**. **Confirmation of the Δ4 mutant genotype by PCR and whole genome sequencing.** Sequencing reads from the Δ4 genome were mapped to the *P. falciparum* reference genome using Bowtie2 (VEuPathDB Galaxy) (Alvarez-Jarreta et al., 2024) and visualized using Integrative Genomics Viewer (IGV) (Robinson et al., 2011). The images captured from IGV confirm the absence of reads within the open reading frames (ORF) of (A) *Pf*MPC1 in chromosome 13, (B) *Pf*MPC2 in chromosome 14, (C) DTC in chromosome in chromosome 8, and (D) YHM2 in chromosome 12. Gene deletion of (E) *Pf*MPC1, (F) *Pf*MPC2, (G) DTC, and (H) YHM2 in the Δ4 mutant was confirmed by PCR amplification (Δ5' and Δ3' products). Intact gene loci (5' and 3' products) were detected only in the NF54$^{attB}$ parental line (blue) and not in the Δ4 line (red). The nucleotide positions of the *Pf*MPC1, *Pf*MPC2, DTC and YHM2 genes in their respective chromosomes, along with CRISPR homology arms HA1 and HA2 and the deleted regions in Δ4 parasites, are listed.

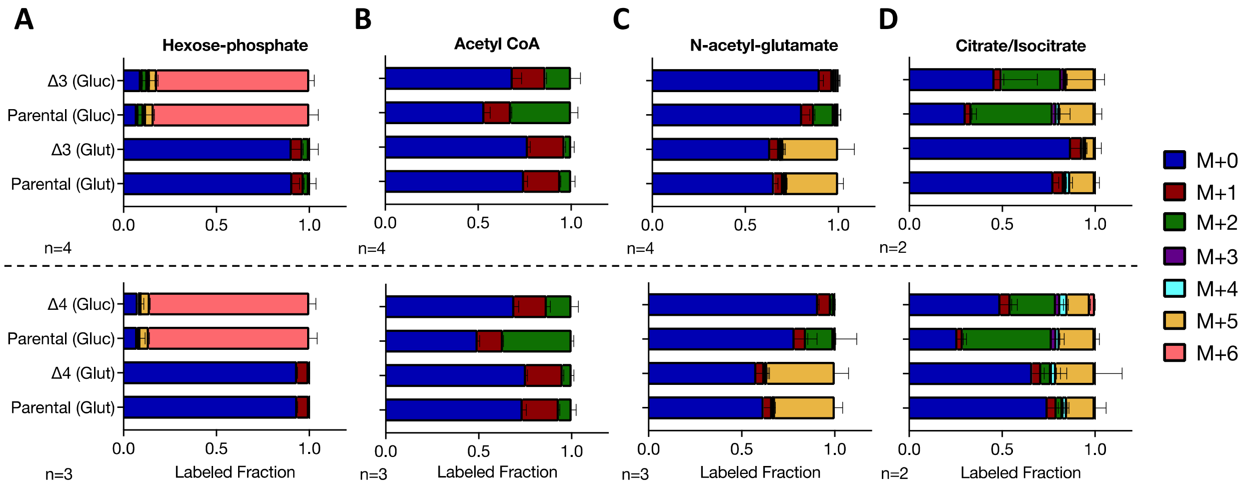


**Fig. S9**. **Deletion of DTC and/or YHM2 does not impact the incorporation of glucose into acetyl-CoA and derivatives.** Fraction of isotopically labeled metabolites when parental, Δ3 or Δ4 parasites are incubated with labeled glucose (Gluc) or glutamine (Glut). Color coding denotes the mass (M) of the parent compound plus the mass shift from the incorporation of heavy carbon isotope ($C^{13}$). Labeling data are expressed as the fraction of the total metabolite pool observed in 'n' independent experiments. Error bars represent the standard deviations from the mean.

| Primers to create p15-SFG plasmid | |
|---|---|
| **Primer Name** | **Sequence (5' → 3')** |
| p15.SFG-F | GGACAACGTACGATGGTATCAAAAGGTGAGGAATTATTTAC |
| p15.SFG-R | CTCGACCTTAAGCCCGGGTCATTTATATAATTCATCCATTCC |
| **Primers to create attB integration lines** | |
| **Primer Name** | **Sequence (5' → 3')** |
| *Pf*MPC1.SFG-F | GGACAACCTAGGATGTCTAATGTAGGATCATTTTTTCATAATG |
| *Pf*MPC1.SFG-R | CACCATCGTACGTTTTGCTGTTATTTTTATTTTTTGATGTTTCG |
| *Pf*MPC2.SFG-F | CGACAACCTAGGATGGAAATAATAAAGAAAATATTTTATCCAAACATCATACC |
| *Pf*MPC2.SFG-R | GCCATCGTACGCCTTTCTTTTTCCTTCATATATACATTATATTTATATTGTC |
| YHM2.SFG-F | TTTTAATTTTTTTTACAACCTAGGATGAGTAATGATTTAATAAATGGTTC |
| YHM2.SFG-R | CCTTTTGATACCATCGTACGAAAATATTTATTTAATATTTTTATGCCAGTAAC |
| **Primers to confirm genotype of attB integration lines** | |
| **Primer Name** | **Sequence (5' → 3')** |
| cg6-F | CTTAACATCCTGTGAAGTTACCCAGGATCC |
| cg6-R | GATACTAATCAAACTGATGAGGAAGTTTTCC |
| attB.int-5' | GCAGTGTGGAATTCCCTGCA |
| attB.int-3' | TTAAGTGTAGTTAATTCATCAAATAGCATGC |
| **Primers to amplify homology arms for gene knockout constructs** | |
| **Primer Name** | **Sequence (5' → 3')** |
| *Pf*MPC1.HA1-F | GCCACGAGCGGCCCGTGTGTATTTTCCTCTGTATTTATAGAGACCC |
| *Pf*MPC1.HA1-R | AAGCGCAGCGGCCCCAATTAGCTAAGGGAGCCCAAAGC |
| *Pf*MPC1.HA2-F | CGACAGACGCCGGGACGACCGTTTTAACAATATATAGTTTATTGTTTATGAG |
| *Pf*MPC1.HA2-R | GGCCACCAGCCGGCCTTCCCTATACGTCCTGTTCATT |
| *Pf*MPC2.HA1-F | GCCACGAGCGGCCATGGAAATAATAAAGAAAATATTTTATCCAAACATCA |
| *Pf*MPC2.HA1-R | AAGCGAGCGGCCGCCCAAAAATGAATAGTCAAGATACCAGTATC |
| *Pf*MPC2.HA2-F | CGACAGACGCCGGCACGATTTGCTTATATGATTAAACCTAGGAACC |
| *Pf*MPC2.HA2-R | GGCCACCAGCCGGGTCCTATCCTAGATATTTGATAAAAAGATGTCATACTCATA |
| DTC.HA1-F | GTGCCACGAGCGGCCGATGGACAGAGATATAGCTAAATATGATTTAG |
| DTC.HA1-R | TAAGCGCAGCGGCCTCCACCAGCTGCTAGAGCAC |
| DTC.HA2-F | TTCGACAGACGCCGGTTCGGTTCCAACTATAGCTAG |
| DTC.HA2-R | ATGGCCACCAGCCGGCTTGTGGTTGTGTAATCAAGTTAAG |
| YHM2.HA1-F | CCGTGCCACGAGCGGCCGCAGAACAATTTTGAAATGAGTAATGAT |
| YHM2.HA1-R | CTTAAGCGCAGCGGCCGCATGCTCCACCAATAAAACCAC |
| YHM2.HA2-F | GAATTCGACAGACGCCGGCTACTGAATGGATAAGAAATATAGTTATGG |
| YHM2.HA2-R | CATGGCCACCAGCCGGCAATCCATATTGTATTATCTACAAATGC |
| **Primers to confirm genotype of knockout parasite lines** | |
| **Primer Name** | **Sequence (5' → 3')** |
| pl8HA1-R | CAAGTATATATTTTGTTTCTATAAATTGATATC |
| pl8HA2-F | CATATTTATTAAATCTAGAATTCGACAGACGCCG |
| *Pf*MPC1.5'-F | CATACATATATGAGAAATCATTTAAAGATTGGG |
| *Pf*MPC1.3'-R | AGTAGTTAAATAATATGATATGTGAGAATGTAAAAC |
| *Pf*MPC1.P.5'-F | GGTTTTGTTATAGCCGGTAGG |
| *Pf*MPC1.P.3'-R | CGGGTTCTTTCTTTAAATCGTTGCATC |
| *Pf*MPC2.5'-F | GTTTGTATTTACCAAAGCACAAAATATAATAATACATAC |
| *Pf*MPC2.3'-R | GAAATAAACTTTTGATGTTCTTGTAATCTCTGCCTTAG |
| *Pf*MPC2.P.5'-F | GGTCCATATCATTAGCCAACATTGC |
| *Pf*MPC2.P.3'-R | GCAATCCAGTTAAACAGATAGCTGTTAAATTC |
| DTC.5'-F | GGAAATATTTCGTTAGATAATATTCCTTACC |
| DTC.3'-R | CAACATTTGTGTTTTCTCAACCCCAC |
| DTC.P.5'-F | GGAAATCCTGCAGATTTATCTTTAATTAG |
| DTC.P.3'-R | GAAATTCTATATAATGCATTAAACACACCAG |
| YHM2.5'-F | CCGTACTGTAATAAACTACTTTTAAC |
| YHM2.3'-R | GCTTATAAGCTATACAATACGAGGTTG |
| YHM2.P.5'-F | GACACCTTGTACATTTTTCATAACTG |
| YHM2.P.3'-R | GTCTTAAACACATAGCAGTGTTCC |

**Table S1**. **List of primers for plasmid construction and genotyping.** Sequences colored in blue depict restriction sites used for ligation-dependent cloning. Sequences colored in red depict the ~15 bp overhang required for In-Fusion cloning.
